# Human cortico-vascular assembloids reveal a CELF2-AHNAK-dependent switch from neuronal to endothelial tropism in glioblastoma cells

**DOI:** 10.1101/2025.10.21.683634

**Authors:** Michele Bertacchi, Annie Ladoux, Alexa Saliou, Gwendoline Maharaux, Clément Peux, Laurent Turchi, Béatrice Polo, Hervé Chneiweiss, Elias El-Habr, Marie Pierre Junier, Fabien Almairac, Fanny Burel-Vandenbos, Michèle Studer, Thierry Virolle

## Abstract

Glioblastoma (GB) remains one of the most aggressive human cancers, driven by profound cellular plasticity and dynamic interactions with the neurovascular niche. Yet, existing preclinical models fail to reproduce the human brain-vascular interface, limiting studies of tumor-host crosstalk and invasion. To overcome this gap, we developed a human induced pluripotent stem cell (hiPSC)-derived cortico-endothelial (CO+EO) assembloid model by fusing cortical and endothelial organoids from the same genetic background. These assembloids spontaneously form branched vascular networks enriched in tight junction proteins (CLDN5, OCLN, ZO-1), thereby recapitulating a key blood-brain barrier (BBB)-like property and providing a physiologically relevant human platform for exploring glioblastoma-neurovascular interactions. Using this system, we uncover a previously unrecognized CELF2-dependent glioblastoma stem cell (GSC) tropism. CELF2-expressing GSCs preferentially infiltrate neural regions, where they trigger neuronal apoptosis and disrupt endothelial tight junction integrity, supporting an aggressive phenotype. Conversely, CELF2-deficient GSCs lose neurotropism, acquire mesenchymal features, and are redirected toward vascular compartments. This endothelial affinity requires the scaffold protein AHNAK, strongly expressed at the plasma membrane of CELF2-deficient cells and enriched at tumor-endothelial interfaces. AHNAK knockdown abolished endothelial infiltration, demonstrating its critical role in vascular tropism. Analysis of patient GB samples confirmed that CELF2-positive tumor cells are enriched in poorly vascularized, mitotically active areas and excluded from vessel-rich zones, closely paralleling assembloid findings. Transcriptomic profiling further revealed that CELF2 promotes a neuronal progenitor-like program while repressing mesenchymal and vascular-associated gene expression, thereby shaping tumor identity, invasive behavior, and tissue preference. Collectively, our study introduces CO+EO assembloids as an original human model of glioblastoma plasticity at the neurovascular interface. We identify CELF2 as a master regulator of GSC tropism and AHNAK as a mediator of vascular affinity, unveiling a novel molecular axis that governs glioblastoma invasion and highlighting new therapeutic opportunities.

**Key points:**

- Developed cortico-endothelial assembloids mimicking the brain microenvironment.
- Identified CELF2 level as a determinant of glioblastoma cell tropism toward neuronal or endothelial niches.
- Showed CELF2-positive GB cells disrupt endothelial tight junctions, unlike CELF2-negative cells.

**Importance of the study:** This study presents a novel, physiologically relevant in vitro model of glioblastoma (GB) that faithfully recapitulates the complex human brain microenvironment, including both vascular and neuronal components. By combining cortical and endothelial organoids derived from human iPSCs into cortico-endothelial assembloids (CO+EOs), the model enables detailed analysis of GB cell behavior, including migration, tissue tropism, and impact on the neurovascular interface. The work identifies CELF2 as a key molecular determinant of GB cell tropism and neurovascular interaction. The discovery that CELF2-expressing GB cells preferentially infiltrate neuronal tissue and disrupt endothelial tight junctions, whereas CELF2-deficient cells favor endothelial regions without compromising tight junction integrity, offers critical insights into tumor heterogeneity and mechanisms of invasiveness.

These findings not only enhance our understanding of how GB cells interact with distinct microenvironmental niches but also provide a powerful platform for testing therapeutic strategies targeting tumor-microenvironment interactions.

**Graphical Abstract:** Schematic representation of CO+EO assembloids comprising an endothelial (yellow) and a cortical (red) domain. On the left, control CELF2-expressing GB cells preferentialy infiltrate the neuronal compartment, where they trigger neuronal apoptosis and disrupt BBB-like tight junctions upon contact with endothelial cells. On the right, CELF2-deprived GB cells exhibit reduced neural invasion and increased tropism for endothelial cells, consistent with a mesenchymal signature shift partially driven by elevated AHNAK expression. These cells exert limited impact on neuronal viability and BBB integrity, and display partial loss of GSC identity, as indicated by reduced OLIG2 expression. The CO+EO assembloid model thus recapitulates key features of cortico-endothelial human tissue heterogeneity and provides a reliable platform to dissect the molecular mechanisms underlying GSC tropism and their tissue-specific interactions

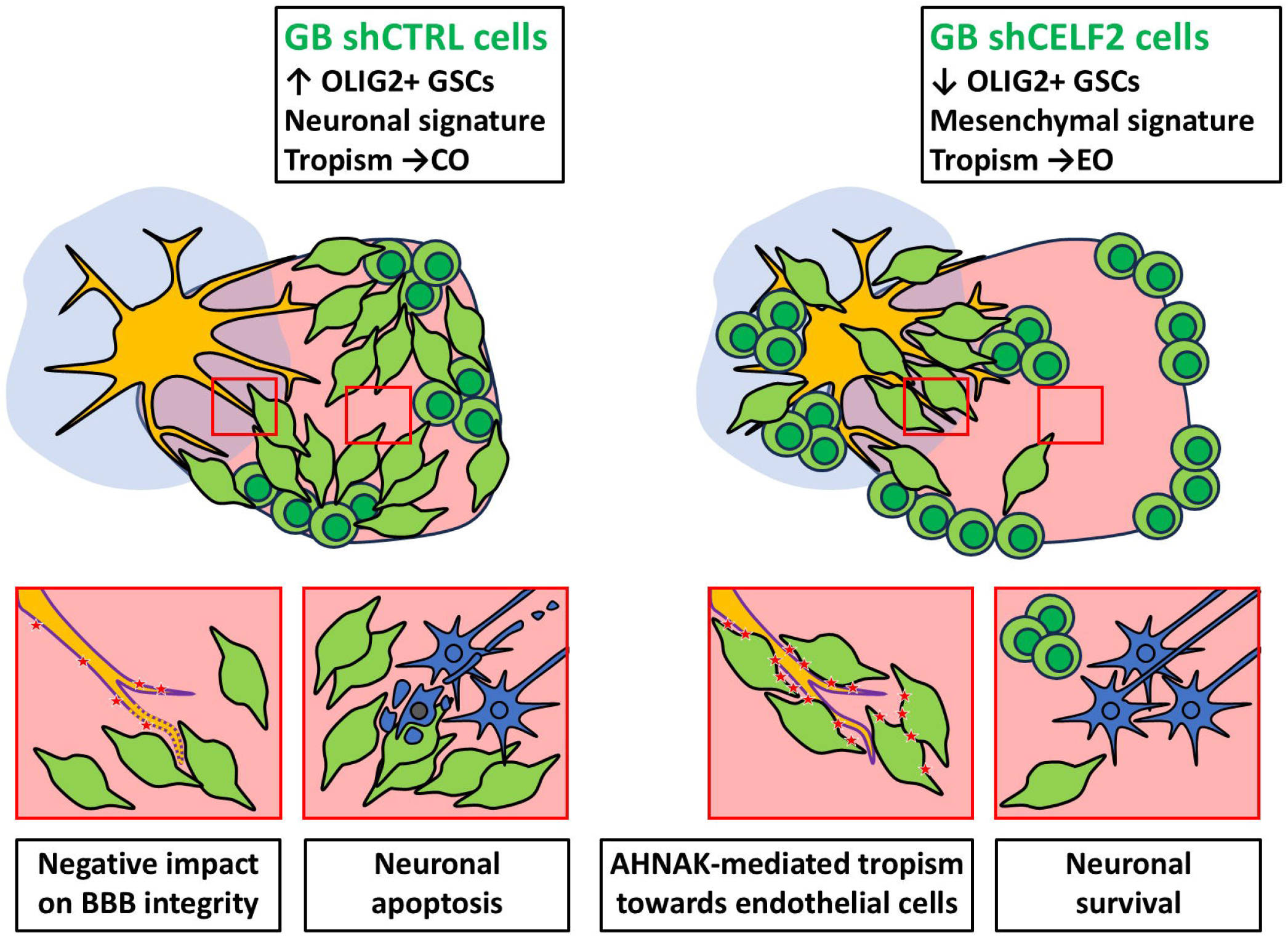

## INTRODUCTION

Glioblastomas (GB), the most common type of primary brain tumor in adults, represent a major public health challenge due to their aggressive nature and poor survival outcomes [1]. Current standard treatments [2] offer only limited improvements in patient life expectancy, largely due to the presence of treatment-resistant cells, the glioma stem cells (GSC), which drive tumor recurrence and progression through their aggressive and resilient characteristics [3, 4]. In close interaction with the tumor microenvironment, GB cells display a high degree of plasticity, enabling reversible transitions between an aggressive proliferative GSC state, characterized by the expression of stem cell markers such as NANOG, SOX2, and OLIG2, and a more differentiated, quiescent, and less aggressive phenotype marked by GFAP expression [5, 6]. This dynamic plasticity underlies the functional hierarchy and intra-tumoral heterogeneity that define GB pathophysiology. Advancing our understanding of key GB cell behaviors, including proliferation, infiltration, tropism, and crosstalk with the surrounding normal brain tissue, is critical for identifying new therapeutic targets and developing more effective therapeutic strategies.

A hallmark of glioblastoma is its characteristic architecture, comprising a dense tumor core, including hypoxic and necrotic areas with increased vascularization, and an infiltrative tumor periphery, each exhibiting distinct transcriptomic signatures [7] [8]. The tumor core typically contains a mix of GSCs and differentiated tumor cells, which may intermingle or segregate into discrete micro-territories, exhibiting either strong or poor level of mitotic activity [6, 9, 10]. The formation of these micro-territories likely results from continuous interactions between tumor cells and the surrounding heterogeneous brain tissue. A major component shaping the tumor core is the microvasculature [7]. The newly formed blood vessels are often dysfunctional and prone to thrombosis, contributing to hypoxic conditions and the emergence of necrotic areas [7]. Concomitantly, the integrity of the blood-brain barrier (BBB), normally maintained by tightly connected endothelial cells, is compromised, allowing abnormal immune cell infiltration and fostering a pro-inflammatory microenvironment [11]. GB cells can migrate along blood vessels to infiltrate the surrounding brain parenchyma [12], contributing to the disruption of BBB, parlty through the loss of tight junctions between endothelial cells [13].

Neurons constitute another critical component of the tumor microenvironment. At the invasive tumor margin, GB cells can interact with neurons, establish synapse-like connections, and integrate into neural circuits, while upregulating genes associated with a neuronal identity [14]. In contrast, more differentiated GB cells exhibit a reduced neuronal gene expression profile [14]. Emerging evidence suggests that this functional and structural integration into neuronal networks is a key driver of tumor progression [15].

These findings highlight the key role of the tumor microenvironment, especially the interactions with endothelial and neuronal cells, for GB homeostasis and progression. Thus, investigating GB cells within a complex and heterogeneous environment is essential to uncover relevant molecular mechanisms that govern their behavior and that can be useful for the design of novel therapeutic strategies targeting tumor cell invasion.

We developed a human cerebral organoid model from induced pluripotent stem cells (hiPSCs) to study glioblastoma (GB) cell interactions, infiltration, and tissue tropism. By combining cortical (COs) and endothelial organoids (EOs) into cortico-endothelial assembloids (CO+EOs), we generated an innovative brain-like tissue with vessel-like structures that mimic key features of the blood-brain barrier (BBB). Co-culture with patient-derived GB cells provides a physiologically relevant system to investigate tumor invasion, migration, morphology, molecular signatures, and interactions with neural and vascular components. We recently showed that the RNA-binding protein CELF2, a member of the CELF/Bruno-like protein family [16], is essential for maintaining GSC markers (NANOG, SOX2, OLIG2), stemness, and tumorigenic potential [10]. In GB samples, CELF2-positive tumor cells were mainly found in proliferative tumor core regions, while CELF2-negative one localized to non-dividing areas [10].

To investigate the role of CELF2 in GB cell infiltration and tropism, we used our CO+EOs model. Our findings reveal that CELF2-expressing and -depleted GB cells showed distinct tropism respectively in neuronal or endothelial compartment. CELF2 expressing cells promote neuronal apoptosis and alters BBB-like tight juncitons. Loss of CELF2 induces the transition to a non-tumorigenic, mesenchymal-like state [17]. We identify AHNAK [18] as a key effector of vascular tropism in CELF2-deficient cells, positioning the CO+EO model as a valuable platform to dissect GB progression and tumor– microenvironment interactions.

## MATERIALS AND METHODS

**(see supplemental material and methods for detailed procedure)**

### hiPSC culture

HMGU1 hiPSCs, derived from neonatal foreskin fibroblasts (ATCC CRL-2522, BJ), were kindly provided by Dr. Drukker (HMGU, Germany) under a Material Transfer Agreement (MTA), and confirmed mycoplasma-free. Cells were maintained on Matrigel-coated plates (Corning, 354234) in mTeSR1 medium (STEMCELL Technologies, #85850) with daily medium changes. Passaging was performed using Versene (Thermo Fisher, 15040066) or, for single-cell dissociation, Accutase (Sigma, A6964). For single-cell culture, 10□µM ROCK inhibitor Y-27632 was added for 24□h post-dissociation to prevent apoptosis.

### Glioblastoma cell line culture

GB5 cells were isolated from primary GBM surgical specimens (University Hospital of Nice), and TG6 cells were kindly provided by Dr. Hervé Chneiweiss (Université Pierre et Marie Curie, Paris). Cells were cultured as previously described [10]. CELF2 knockdown (shCELF2) and control (shCTRL) lines were generated as previously described [10].

### hiPSC and Glioblastoma cell line quality controls

HMGU1 and glioma stem cells were tested annually for mycoplasma contamination by PCR using Myco_F1 and Myco_R1 primers on supernatant collected after 48 hours of culture. Genetic and chromosomal integrity of HMGU1 was evaluated with the hPSC Genetic Analysis Kit (Stem Cell Technologies, #07550), using WT-T12 and PGP1 hiPSC lines as controls. Genomic DNA was extracted from Proteinase K-lysed cell pellets, followed by isopropanol precipitation and ethanol washing. Due to recurrent mutations and chromosomal abnormalities after extended culture, HMGU1 cells were not used beyond passage 26. Pluripotency was confirmed via OCT4 immunostaining prior to downstream applications.

### Neural differentiation of cortical organoids (COs)

HMGU1 cells (25–50k/well) were seeded on Matrigel-coated 24-well plates in mTeSR1/+ until ∼90% confluency, then switched to neural progenitor patterning medium (NPPM) with BMP, TGFβ, and WNT inhibitors (LDN-193189, SB-431542, XAV-939).

At day 7–8, cells were dissociated (Accutase), seeded into 96-well U-bottom plates (60k/well) in NPPM + ROCK inhibitor, and centrifuged to form aggregates. By day 8–9, aggregates were embedded in Matrigel droplets and cultured in bioreactors with NPPM + XAV-939 (LDN and SB removed from day 9– 10).

At day 20, medium was switched to Neural Differentiation Medium (NDM), then to Long-Term Survival Medium (LTSM) by day 30–35. Medium was refreshed every 3–4 days. From day 50–60, large organoids were sectioned monthly to prevent necrosis.

### Mesoderm differentiation for endothelial organoid (EO) production

Mesodermal differentiation was adapted from Wimmer et al. [19]. Dissociated hiPSCs were seeded into ultra-low attachment 96-well plates (4,000 cells/well) in mTeSR1 medium with 5□µM Y-27632 (Tocris) and centrifuged at 125□×□g for 4□min to promote embryoid body (EB) formation. After 2 days, the medium was replaced with DMEM/F12 supplemented with N2/B27 (with vitamin A), 10□µM CHIR99021 (Tocris), and 30□ng/mL BMP4 (ThermoFisher) for mesoderm induction. On day 5, EBs were transferred to N2/B27 medium containing 40□ng/mL VEGF-A and 2□µM forskolin (Tocris) for 2 additional days. For vascular organoid generation, 10–15 mesodermal EBs were embedded in Collagen I/Matrigel (Corning) matrices in 12-or 24-well plates. StemPro34 SFM (ThermoFisher) with 10% FBS, 40□ng/mL VEGF-A, and 20□ng/mL FGF-2 was added to induce vessel sprouting, typically observed within 2–3 days. Medium was partially changed every other day. EOs were harvested after day 10 using modified pipette tips and gentle mechanical dissociation, and used for assembloid formation upon confirmation of vascular sprouting.

### Cortico-endothelial assembloid (CO+EOs) formation

Cortical (CO) and endothelial organoids (EO) were co-cultured in low-adhesion 96-well plates in LTSM + VEGF-A (40□ng/mL) and FGF2 (20□ng/mL). Medium was partially refreshed daily. After 1 week, 10,000 GB cells were added per well, followed by brief centrifugation (300□rpm, 1□min) and 24□h incubation. Assembloids were then transferred to an orbital shaker for continued culture.

### Immunostaining on cryostat sections

Organoids and assembloids were fixed in 4% PFA (4□°C, 4□h), cryoprotected in 1–25% sucrose, embedded in OCT, and cryosectioned (12□µm). Sections were dried, washed with PBS, and antigen retrieval was performed (0.1□M sodium citrate, 95□°C, 10□min). Section were blocked (PBS + 5% serum + 0.3% Triton X-100) and stained using primary antibodies (≥4□h RT or overnight at 4□°C), Alexa Fluor secondaries (1:500), and Hoechst (1:1,000). Imaging was performed with a Vectra Polaris, Zeiss Apotome, or LSM710 confocal microscopes, with analysis in HALO, Fiji, or Photoshop.

### Immunostaining image analysis on HALO software

Tissue segmentation was performed using the Random Forest Classifier in HALO. The AI was trained on marker expression to identify cortex (MAP2/TUJ1), tumor (GFP), and endothelium (CD31+/MAP2–). Once trained, the classifier was applied to additional sections. HighPlex FL modules (v4.2.3/4.2.14) were used for marker-positive cell detection and further analysis.

### Immunostaining and tissue clearing of whole organoids

Fixed organoids were permeabilized (0.5% CHAPS, 3□h, 37□°C), blocked overnight (PBS-azide + 0.3% Triton X-100 + 5% NBCS), and incubated with primary (1:250, 2–3□days) and secondary (1:500, 24–48□h) antibodies. After washes, organoids were cleared in AKS solution (DMSO, TDE, sorbitol, Tris base) for ≥24□h, and embedded in 0.8–0.9% agarose in AKS for 3D imaging via Zeiss Lightsheet 7. Analysis was done in Imaris (v9.6).

### Immunostaining of Cultured Cells

Cells on coverslips were labeled with primary antibodies (see Supplementary Materials), followed by Alexa Fluor-conjugated secondary antibodies (1:1000). Controls without primary or with non-specific antibodies were included. Imaging was done using a Zeiss AxioObserver microscope or Vectra Polaris scanner.

### Western Blot

Proteins were extracted from cells/organoids, separated by SDS-PAGE, and transferred to PVDF membranes. Detection was performed with HRP-conjugated secondary antibodies and ECL reagents. Signals were imaged (Fusion FX7) and quantified using FIJI.

### RT-qPCR

RNA was extracted (TRI-Reagent), reverse-transcribed (MMLV), and analyzed by SYBR Green-based real-time PCR. Gene expression was normalized to 36B4 using the ΔΔCt method. Primer sequences are listed in Table 2.

### Statistical analysis

Data were analyzed using Excel, GraphPad Prism, or BiostaTGV, with results expressed as mean□±□SEM. Cell counts were done manually or via AI (HALO). Statistical tests included Mann– Whitney, t-tests, and ANOVA (p<0.05).

### AHNAK Knockdown

GB cells were transfected with shRNA Clone sets of three constructs against Human AHNAK nucleoprotein in psi-LVRH1H, including a scrambled control clone (ref # CS-HSH111706-LVRH, Labomics S.A., Nivelles, Belgium) using the jet Prime transfection kit (Polyplus-transfection S.A., Illkirch, France) following the recommendations of the manufacturer. Transfected cells were selected and further grown with hygromycin (50µg/ml). The absence of AHNAK expression was analyzed by western blots as previously described.

### RNA-seq Analysis

Bulk RNA-seq from shCTRL and shCELF2 GB5 cells [10] was analyzed in R (ggplot2); differential gene expression and volcano plots were generated using Neftel et al. signatures [17].

## RESULTS

### hiPSC-derived CO+EO assembloids as an *in vitro* model of highly vascularized brain tissue

To model the human brain–vascular interface, we generated cortico-endothelial assembloids (CO+EOs) by fusing 3D cortical organoids (COs) and endothelial organoids (EOs) derived from the same hiPSC line (Fig. 1A). COs were produced using our telencephalic-inducing protocol [20]. To generate Eos via mesodermal induction, we have adapted a previously described protocol [19] (Suppl. Fig. 1). Early mesodermal induction in EOs was confirmed by transient expression of the mesodermal marker Brachyury (BRY). EOs formed within two weeks capillary-like networks of elongated cells expressing endothelial markers (CD31, VE-CADHERIN) and VEGF receptors, suggestive of blood vessel-like structures (Suppl. Fig. 1A,B). VEGF receptor expression (VEGFR1, VEGFR2, NRP1) increased during differentiation, peaking at maturation, consistent with endothelial commitment. Immunostaining confirmed robust endothelial maturation (CD31+, VE-CADHERIN+) and absence of neural contamination (MAP2) (Supp. Fig. 1D-F), indicating lineage specificity.

**Figure 1.**
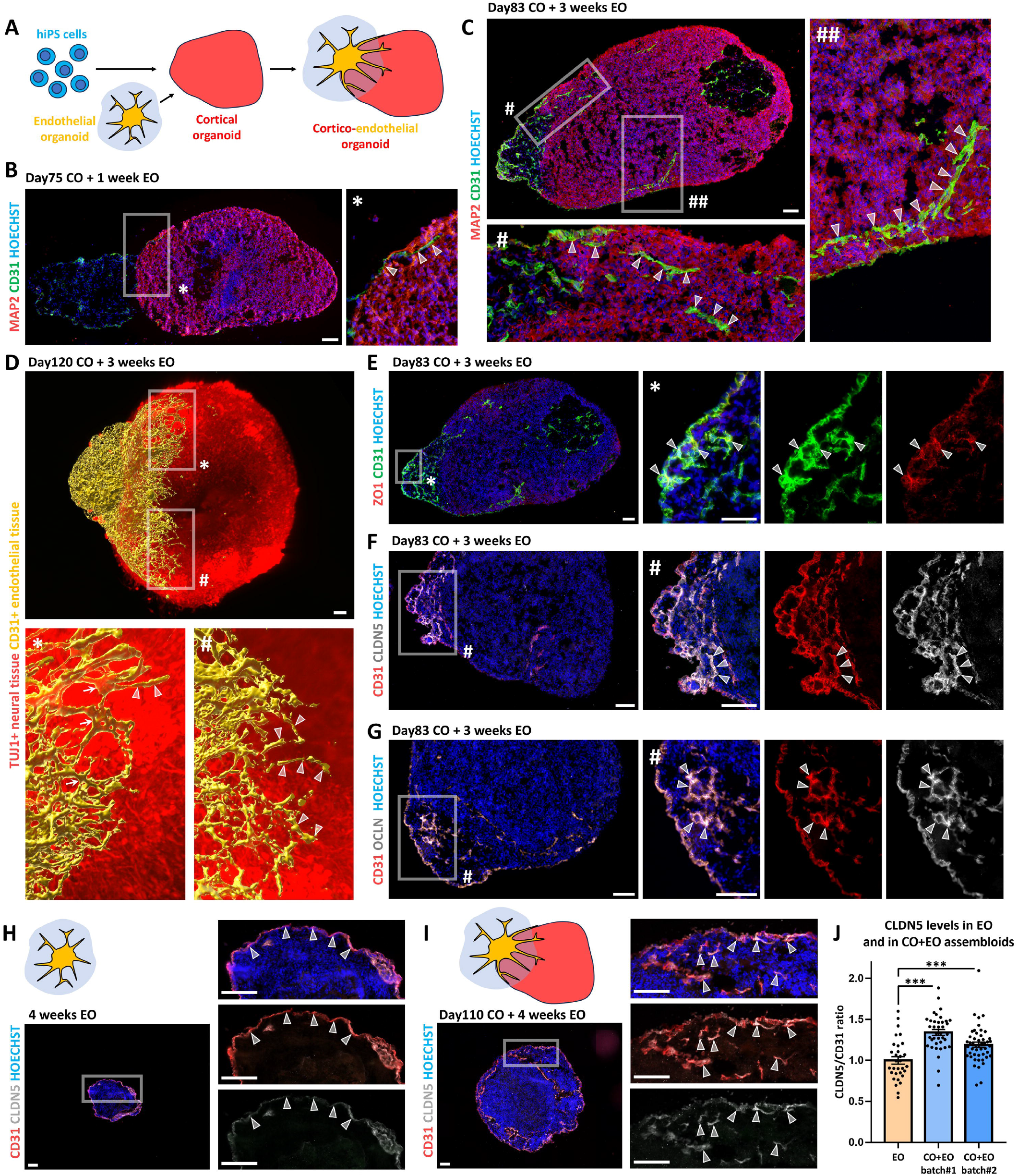
Cortico-endothelial assembloids develop blood vessel-like structures expressing BBB markers. **A)** Schematic representation of the strategy used to fuse cortical organoids (COs; red) with endothelial organoids (EOs; yellow), generating cortico-endothelial assembloids (CO+EOs). **B, C)** MAP2 (red; neural tissue) and CD31 (green; endothelial cells) immunostaining of CO+EOs 1 week and 3 weeks after fusion, as indicated. Arrowheads point to CD31+ cells forming elongated tubular structures within the neural tissue. **D)** 3D reconstruction of the endothelial network (yellow) developing into neural tissue (red) in a CO+EO assembloid after tissue clearing and CD31/TUJ1 immunostaining. Arrowheads point to sprouting endothelial cells, while white arrows indicate branching points in complex ramifications. **E-G)** Co-expression of tight junction markers ZO1 (red in E), CLDN5 (white in F) and OCLN (white in G) with the endothelial cell marker CD31 (green in E and red in F and G); arrowheads highlight double-labeled cells in CO+EOs. **H-J)** Double immunostaining for CD31 (red) and CLDN5 (white) in EOs (H) and CO+EO assembloids (I); white arrowheads in high-magnification images point to double-labeled blood vessel-like structures. Graph in J shows quantification of CLDN5 levels in endothelial cells, measured as a ratio of CLDN5 pixel intensity over CD31 signal, from one EO batch and two independent CO+EO batches. Scale bars: 100 µm.

To generate cortico-endothelial assembloids (CO+EOs), we fused two-month-old or mature cortical organoids (COs) with two-week-old endothelial organoids (EOs) (Fig.1A). Within one week of co-culture, endothelial cells began migrating along the CO surface (Fig. 1B), and by three weeks, they had efficiently infiltrated the organoid parenchyma (Fig. 1C), forming a dense, branched CD31+ network resembling blood vessel-like structures (Fig. 1D). 3D reconstruction of tissue-cleared assembloids revealed extensive vascular branched structures integration throughout the cortical zone (Fig. 1D). To assess blood–brain barrier (BBB) features, we examined tight junction markers in the endothelial compartment. CO+EOs exhibited strong expression of BBB-associated proteins Zona Occludens-1 (ZO-1), Claudin-5 (CLDN5), and Occludin (OCLN) (Fig. 1 E-G). Notably, CLDN5 levels were significantly higher in CO+EOs than in EOs alone (Fig. 1 H-J), indicating that neural-endothelial interactions promote tight junction formation and BBB-like features. These findings demonstrate that CO+EO assembloids spontaneously develop vascularized, brain-like tissue with BBB characteristics, providing a robust human model for studying neurovascular interactions and BBB development in vitro.

These results establish CO+EO assembloids as a robust human model to study brain–vascular interactions and BBB formation.

### CELF2-dependent tropism of glioblastoma cells in CO+EO assembloids

Given the established role of CELF2 in regulating GSC aggressiveness [10], we investigated its influence on GSC infiltration within a complex tissue environment. To this end, we co-cultured previously characterized patient-derived CELF2-expressing GSCs (GB5 shCTRL) and CELF2-depleted counterparts (GB5 shCELF2) with cortico-endothelial assembloids (CO+EOs) (Fig 2A). After three weeks, immunostaining confirmed CELF2 knockdown in GB5 shCELF2 cells (Suppl. Fig. 2A). We then segmented the assembloid surface into neural and endothelial regions and quantified tumor cell distribution (Fig 2B, C). CELF2-expressing cells preferentially localized to TUJ1+ neuronal zones, while CELF2-deficient cells accumulated in endothelial regions (Fig 2B–C; Suppl. Fig. 2B–C). This spatial divergence correlated with a loss of MAP2+/TUJ1+ areas in organoids invaded by shCTRL cells and a reduction in tumor-free endothelial zones in organoids invaded by shCELF2 cells.

**Figure 2.**
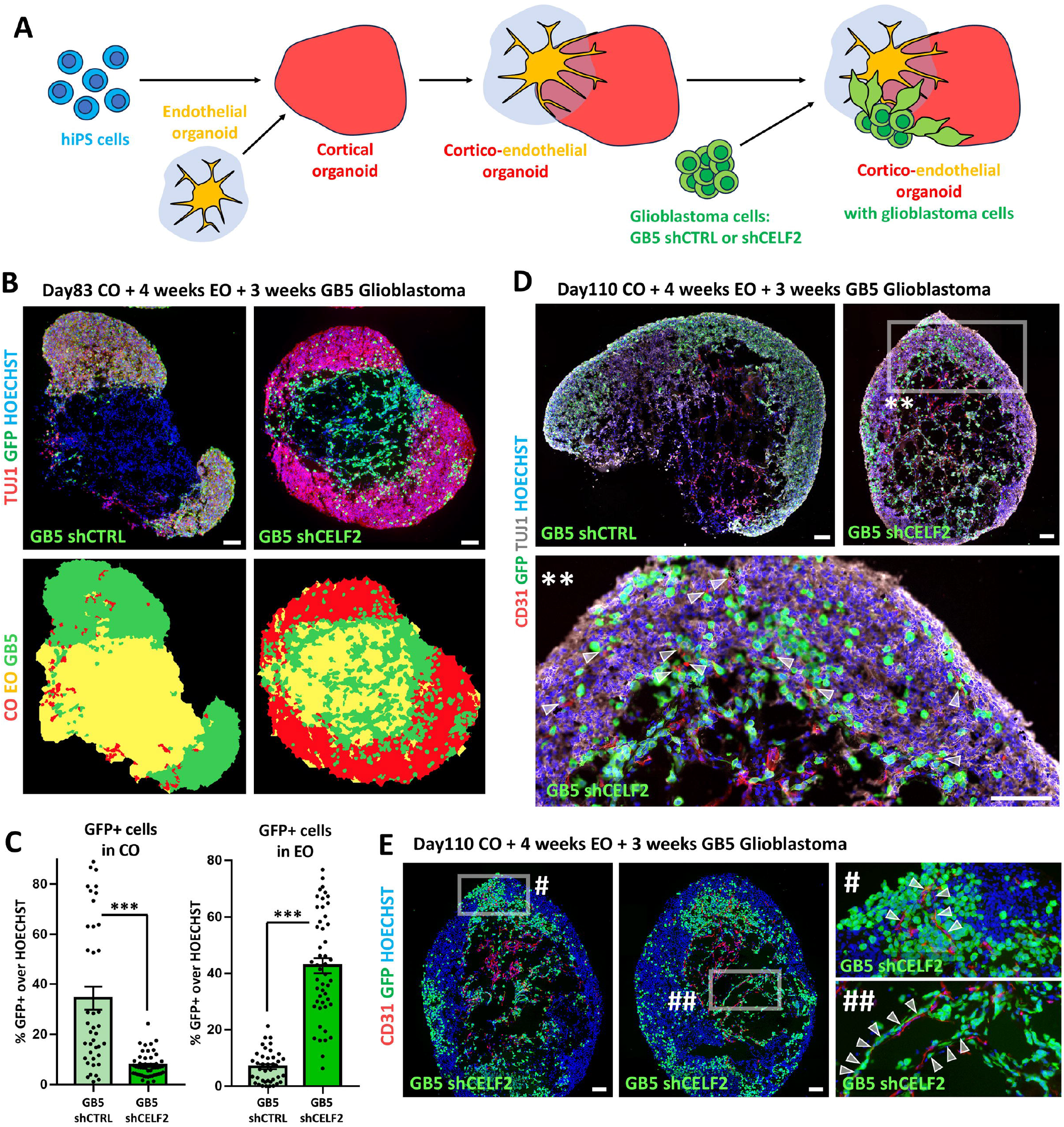
CELF2-dependent tropism of GB cells in cortico-endothelial human assembloids (CO+EOs). **A)** Schematic representation of the co-culture strategy generating cortico-endothelial assembloids (CO+EOs) with GB5 shCTRL and shCELF2 cells. **B**,**C)** TUJ1 (red; neural tissue) and GFP (green; GB5 cells) immunostaining of CO+EOs, 4 weeks after fusion between EOs and COs and 3 weeks post-addition of GB cells. Automated recognition of tumor (green), cortical (red), or endothelial (yellow) surfaces is shown in B, lower row. Graphs in C display the percentages of GFP+ GB cells found in each surface type in human assembloids 3 weeks after GB invasion, with control GB5 shCTRL cells colonizing the cortical domain and CELF2-deprived shCELF2 GB5 cells preferring the endothelial part. **D)** Triple immunostaining for CD31 (red), GFP (green), and TUJ1 (white) of CO+EOs after 3 weeks of co-culture with GB5 shCTRL or shCELF2 cells, as indicated; arrowheads point to CELF2-deprived GFP+ cells in close proximity to CD31+ blood vessel-like structures. **E)** CD31 (red; endothelial cells) and GFP (green; tumor cells) immunostaining of CO+EOs after 3 weeks of co-culture with GB5 shCTRL or shCELF2 cells, as indicated. Arrowheads indicate shCELF2 GB5 cells entering the cortical tissue by migrating along CD31+ endothelial cells (#) or elongating along them (##). Scale bars: 100 µm.

Interestingly, CELF2-deficient GSCs often aligned with CD31+ structures, even when present in neural regions, suggesting that endothelial cells may guide their invasion (Fig 2D–E). These cells also displayed an elongated, migratory-like morphology when interacting with vasculature (Fig 2E). We repeated these experiments using a second patient-derived line, TG6, with or without CELF2 knockdown. Consistent with GB5 findings, TG6 shCTRL cells favored neural zones, whereas TG6 shCELF2 cells were redirected to endothelial compartments (Suppl. Fig. 2D–F). Notably, proliferation (KI67) was unaffected by CELF2 status, but OLIG2+ mitotic cells were significantly reduced in CELF2-deficient populations (Suppl. Fig. 2F–G), confirming a shift toward an OLIG2-negative subpopulation [10] and suggesting that cell proliferation is maintained by an OLIG2-negative subpopulation.

Overall, these results reveal a CELF2-dependent tissue tropism: CELF2-expressing GSCs target neural tissue, while CELF2-deficient cells are redirected toward vasculature. This highlights the CO+EO assembloid as a powerful model for dissecting molecular mechanisms underlying glioblastoma infiltration and tropism.

### Human COs reveal CELF2-dependent dynamics of GB invasiveness in neural tissue

To specifically assess the influence of CELF2-dependent invasive potential of GB cells in a neural-specific context, while avoiding confounding factors introduced by the presence of non-neural tissues, we co-cultured GB5 cells with simple COs (Suppl. Fig. 3A). One week after co-culture, control CELF2-expressing GSCs readily infiltrated the cerebral organoid, whereas CELF2-depleted GSCs remained largely confined to the organoid surface (Suppl. Fig. 3B-E), resulting in reduced surface occupancy in COs (Suppl. Fig. 3E). Morphologically, while CELF2-deficient cells displayed a rounded, non-migratory phenotype within the neural environment, consistent with their reduced invasive potential and in contrast to their elongated, migratory morphology observed during endothelial invasion, the CELF2-expressing counterpart retained an elongated shape, characteristic of actively migrating cells (Suppl. Fig. 3B–D).To directly assess GSC infiltration dynamics, we quantified the spatial distribution of tumor cells across the organoid depth. By two weeks, CELF2-expressing cells had migrated into the organoid core, while CELF2-depleted GCSs remained largely confined to the surface or peripheral regions (Suppl. Fig. 3F,G). These differences became even more pronounced at three weeks, with CELF2-expressing GB5 cells forming extensive, proliferative aggregates deep within the cortical tissue, while CELF2-deficient cells formed smaller, restricted clusters at the periphery (Suppl. Fig. 3H), resulting in reduced tumor area occupancy (Suppl. Fig. 3I). Similar results were obtained with the TG6 GB cell line (Suppl. Fig. 4). Light sheet microscopy of tissue-cleared COs confirmed that CELF2-expressing tumor cells penetrated the organoid core, whereas CELF2-depleted cells remained predominantly localized at the outer edge (Suppl. Fig. 3J). Consistent with findings from CO+EO assembloids (Suppl. Fig. 2F, G), the CELF2-depleted cell population co-cultured with COs alone, retained proliferative capacity despite their reduced invasiveness and lower abundance of OLIG2+ mitotic cells (Suppl. Fig. 3K, L). Collectively, these findings demonstrate that CELF2 directly influence GSC invasiveness, migration, proliferation, and stemness, ultimately impacting GB invasive potential in human brain-like tissue.

### CELF2-dependent GB tropism toward human endothelial tissue

To further validate the endothelial tropism of GB cells driven by CELF2 depletion, initially observed in CO+EO assembloids, we investigated GB behavior in an endothelial-only context by co-culturing CELF2-expressing (shCTRL) and CELF2-depleted (shCELF2) GB5 or TG6 cells with endothelial organoids (EOs) alone (Suppl. Fig. 5A). The presence of mature endothelial cells within the EOs was confirmed by CD31 immunostaining after four weeks of differentiation (Suppl. Fig. 5B). GB cells were subsequently introduced and co-cultured was maintained with the EOs for an additional three weeks to allow their integration and assess their capacity for adhesion and infiltration. In this simplified setting, shCELF2 cells exhibited enhanced integration into the capillary-like vascular structure, adhering to and infiltrating the endothelial network (Suppl. Fig. 5C). In contrast, shCTRL cells largely failed to penetrate the EOs and instead formed compact aggregates along the organoid periphery (Suppl. Fig. 5C), displaying a distinct behavior from that observed in COs or CO+EOs. To quantify these differences, we employed AI-based image analysis (HALO software), to define the endothelial region within the EOs and assess the spatial distribution of GB cells (Suppl. Fig. 5D). This analysis revealed a significantly higher number of CELF2-depleted cells located within the endothelial region for both GB5 and TG6 cell lines, compared to their CELF2-expressing counterparts (Suppl. Fig. 5E, F). These findings underscore the role of CELF2 in directing GB cell tropism toward neural rather than vascular regions, revealing distinct tissue- and CELF2-dependent patterns of invasion.

### Patient tumor analysis confirms CELF2-dependent GB cell tropisms identified in CO+EO assembloid models

We next examined the pathophysiological relevance of CELF2-dependent tropism observed in CO+EO assembloids by analyzing human GB tissue samples. As expected, given the GB intra-tumoral heterogeneity, non-mitotic tumor regions often adjacent to necrotic zones, showed strong immunoreactivity for the astrocyte differentiation marker GFAP and were frequently associated with abnormal dense vasculature (Suppl. Fig. 6). In contrast, mitotic tumor regions were characterized by sparse vascularization (Suppl. Fig. 6). To assess the spatial distribution of CELF2-positive GB cells, we performed immunohistochemical staining on tumor sections from five different GB patients using a CELF2-specific antibody, and quantified the proportion of CELF2-positive cells in highly versus poorly vascularized regions (Fig. 3). Strikingly, CELF2-positive cells were significantly enriched in poorly vascularized areas and markedly under-represented in vessel-rich regions (Fig 3 and Suppl. Fig. 7). This inverse correlation between CELF2 expression and vascular density was consistent and statistically significant across all patient samples analyzed. These in vivo findings closely parallel our in vitro observations in CO+EO assembloids, where CELF2-expressing GSC preferentially localized to neural compartments rather than endothelial zones. Together, these data establish a strong correlation between the CO+EO model and patient tumor biology, underscoring the pathophysiological relevance of the CO+EO assembloid model for dissecting the molecular and spatial determinants of GB cell tropism.

**Figure 3.**
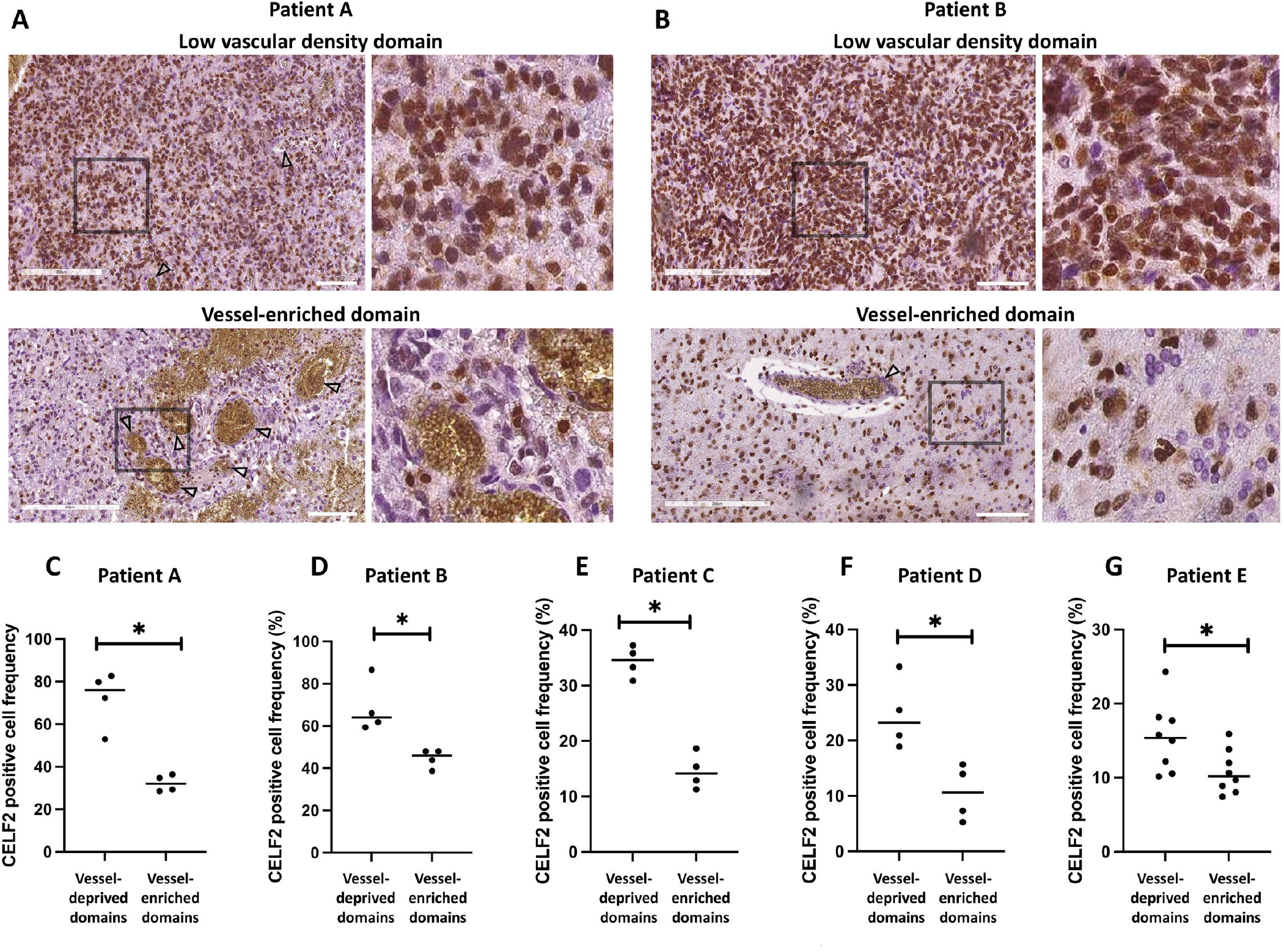
CELF2+ cell percentage in vessel-enriched and vessel-deprived areas of patient GB samples. **A, B)** Exemplary immunohistochemistry stainings of GB samples from two patients, showing CELF2 (brown) over hematoxylin-eosin staining that highlights nuclei (violet) in areas with few and small blood vessels (low vascular density domains; upper row) and in highly vascularized areas with large and numerous blood vessels (vessel-enriched domains; bottom row). Arrowheads highlight blood vessels. CELF2 immunohistochemistry stainings on other 3 GB patients are shown in Suppl. Figure 6. **C-G)** Graphs showing the percentages of CELF2+ cells in vessel-deprived regions and in vessel-enriched regions within the GB tumor sections of five different patients. Scale bars: 100 µm.

### CELF2-dependent impact on neuronal and endothelial components of CO+EO assembloids

We next investigated the impact of patient-derived GB cell invasion on MAP2-positive neurons and endothelial cells within the CO+EO host tissue, and whether these effects depend on CELF2 expression (Fig. 4A). To this aim, we first evaluated neuronal integrity by co-staining for MAP2 and the apoptotic marker cleaved Caspase-3 (Fig. 4B,C). We observed increased neuronal apoptosis in CO+EOs co-cultured with highly invasive, CELF2-expressing GB5 or TG6 control cells compared to assembloids co-cultured with CELF2-depleted counterparts (Fig. 4B,C). Given that CO+EO assembloids support endothelial maturation, as evidenced by increased expression levels of tight junction proteins (Fig. 1H-J), we next investigated whether GB cell infiltration could compromise the integrity of this BBB-like structure. To assess the structural integrity of the endothelial barrier, we quantified the ratio of CLDN5 (a tight junction marker) to CD31 (an endothelial marker) in regions proximal or distal to GB cells. No significant changes in the CLDN5/CD31 ratio were observed in zones distant from tumor cells or in any condition involving CELF2-deficient cells. However, in regions where CELF2-expressing GB cells were in direct contact with endothelial cells, the CLDN5/CD31 ratio was significantly reduced (Fig. 4E,F), suggesting a loss of endothelial barrier integrity. Together, these results indicate that aggressive CELF2-positive GSCs exert detrimental effects on the CO+EO microenvironment by inducing neuronal apoptosis and disrupting BBB-like endothelial structures. In contrast, CELF2 depletion, previously shown to convert aggressive GSCs into a less aggressive phenotype [10], is sufficient to prevent these deleterious effects and preserve host tissue integrity.

**Figure 4.**
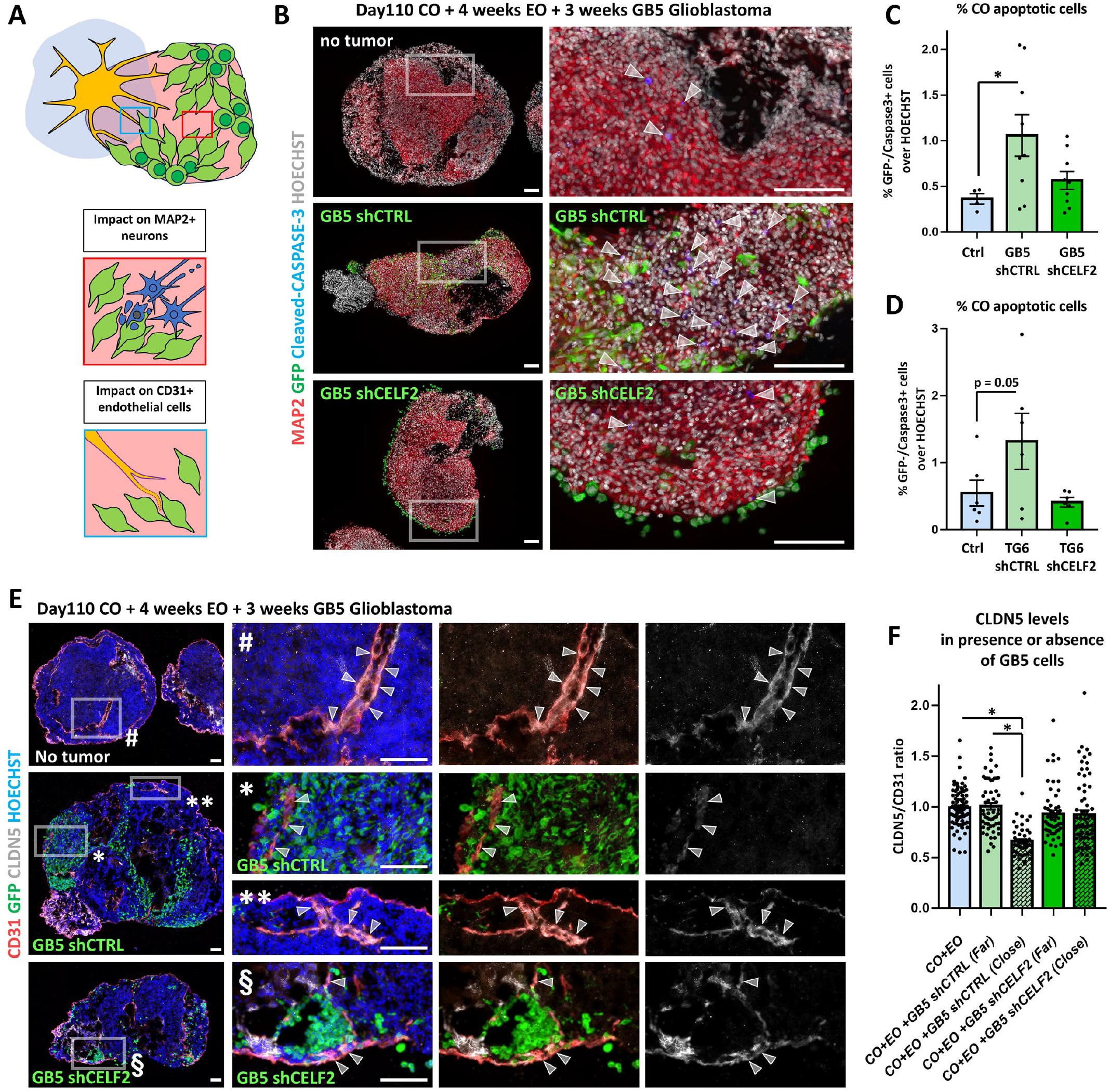
Neuronal survival and BBB integrity in CO+EOs upon GB cell invasion. **A)** Schematic representation of a CO+EO assembloid invaded by GB cells (green); MAP2+ neurons and CD31+ endothelial cells are the main cellular populations interacting with GB cells evaluated here. **B-D)** Triple immunostaining for MAP2 (red; neural tissue), GFP (green; GB cells), and cleaved CASPASE3 (blue; apoptotic cells) in control (top row in B), GB5 shCTRL-invaded (middle row), and GB5 shCELF2-invaded (bottom row in B) CO+EOs. Graphs show the percentage of apoptotic neuronal cells (double MAP2+ and CASPASE3+) in control assembloids and in assembloids invaded by shCTRL or shCELF2 GB5 (graph C) or TG6 (graph D) cells. **E**,**F)** Triple immunostaining for GFP (green), CD31 (red), and CLDN5 (white) in CO+EO assembloids after 3 weeks of co-culture with shCTRL or shCELF2 GB5 cells; pixel intensity of CLDN5 in CD31+ cells under different conditions (shCTRL versus shCELF2 cells; far versus close to GB cells) is displayed in graph F, showing decreased CLDN5 levels when control CELF2-expressing cells come in close contact with endothelial cells. Scale bars: 100 µm.

### Molecular signatures correlating with CELF2-dependent GSC tropism in CO+EO assembloids

To further understand how CELF2 expression influences GSCs phenotype and behavior within CO+EO assembloids at the molecular level, we re-analyzed bulk RNA sequencing data previously generated from GB5 cells with or without CELF2 expression [10]. Specifically, we examined gene expression patterns associated with the neural, astrocytic, oligodendrocyte progenitor, and mesenchymal GB subtypes, as defined by Nelftel et al. [17]. CELF2-expressing GB5 cells displayed a robust neuronal transcriptional signature, including genes involved in neuronal development (*DCX, STMN2, TAGLN3, CRABP1, DLL3*), GB progression (*NEU4, BCAN1*), epilepsy (*NXPH1*) and hypoxia (*SDH*) (Fig. 5A). In contrast, CELF2-deficient GB5 cells exhibited a transcriptional profile consistent with a mesenchymal identity, characterized by CHI3L1 (a key mesenchymal identity driver [21]), *EFEMP1* (a tumor suppressor regulator [22] [23]), *NNMT* (promoting mesenchymal GB proliferation [24]) and *SERPING1* (linked to poor GB prognosis [25]) expressions (Fig. 5A) (supplemental table 3).

**Figure 5.**
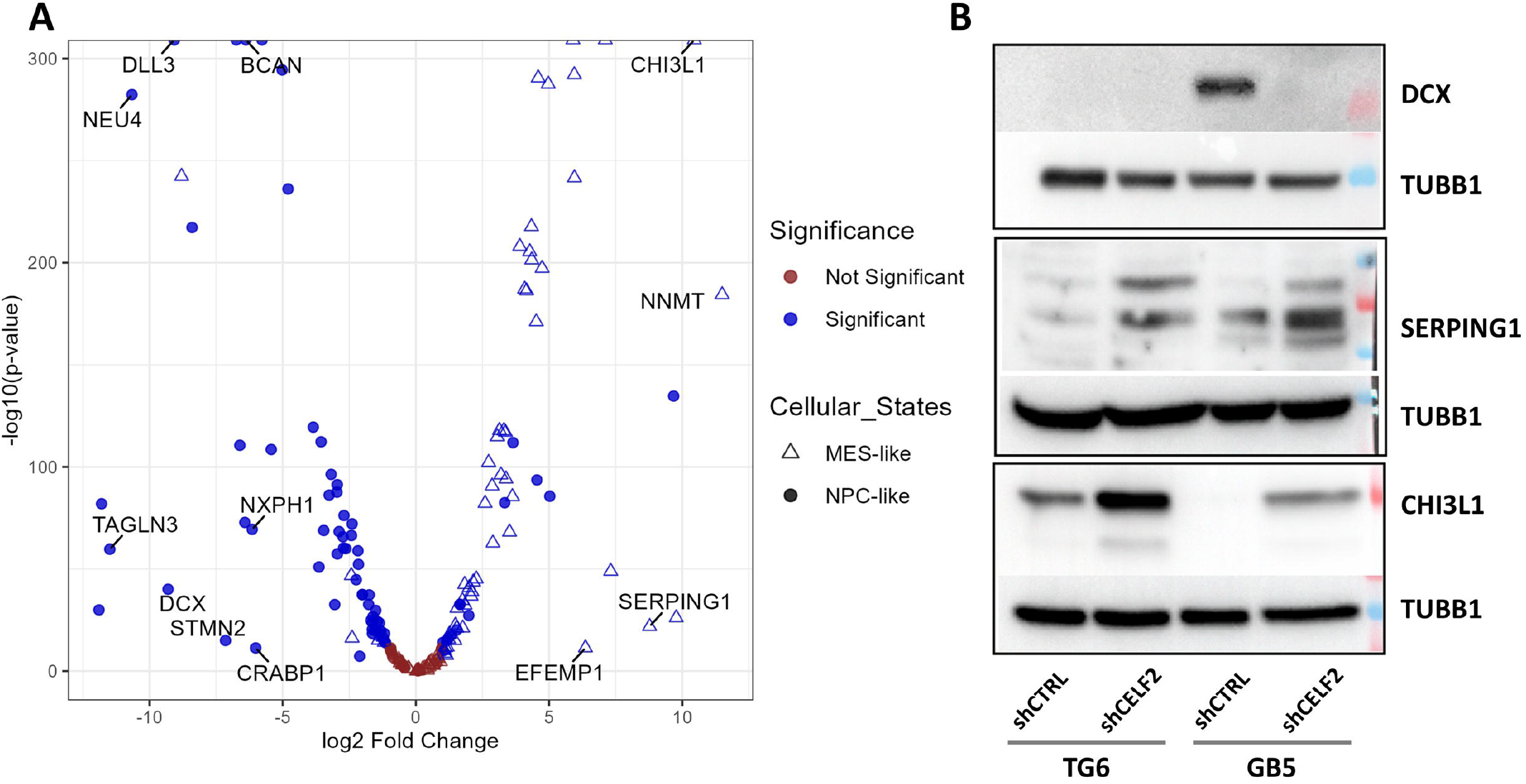
CELF2-dependent molecular signatures in GB cells. **A)** Volcano plot showing upregulated and downregulated genes (x-axis: fold change; y-axis: corresponding p-value) comparing GB5 shCELF2 and GB5 shCTRL cells, highlighting genes affected by CELF2 suppression and the resulting neuronal or mesenchymal signatures. **B)** Western Blot analysis for the protein expression of representative genes belonging to the neural (DCX) or mesenchymal (SERPING1, CHI3L1) signatures in GB5 and TG6 cells expressing or not CELF2.

These transcriptional differences were corroborated at the protein level using representative neural and mesenchymal markers (Fig 5B). Comparison with transcriptional datasets from Ren et al [26] further supported the existence of two distinct CELF2-dependent transcriptional programs: one favoring a neuronal-like identity and the other a mesenchymal one. Genes enriched in CELF2-expressing cells were associated with synapse organization and positive regulation of neural development and neurogenesis, including *NEUROD1, STMN2, SEMA5A, PCP4, NLGN3*, and *BCAN* (Supplementary Table 1). Conversely, CELF2-deficient cells upregulated genes associated with mesenchymal transformation and vascular interaction, including CHI3L1, HSPG2 (a vascular extracellular matrix component), *VEGFA* and *ADM* (both involved in vascular development and permeability) (Supplementary Table 1). Notably, several integrin genes were also upregulated in CELF2-deficient cells, potentially contributing to their enhanced adhesion and endothelial tropism. Together, these findings indicate that CELF2 promotes a neural progenitor-like transcriptional state while repressing mesenchymal and vascular-associated programs, thereby shaping the distinct invasive behavior and microenvironmental preferences of GB cells in a tissue-specific manner.

### AHNAK contributes to the endothelial tropism in CELF2-deficient GB cells

Among the genes upregulated within the mesenchymal transcriptional program of CELF2-deficient GB cells, we focused on the large scaffold protein AHNAK to assess its potential role in mediating endothelial affinity in the CO+EO assembloid model (Figure 6A). Western blot analyses of both GB5 and TG6 cell lines confirmed that AHNAK protein was robustly expressed in CELF2-deficient GSCs, while its expression was weak in CELF2-expressing cells and endothelial organoids (Fig. 6B). On the other hand, immunofluorescence staining revealed high levels of AHNAK in GB5 shCELF2 cells, with prominent localization at the plasma membrane (Fig. 6C). Confocal imaging of CO+EO assembloids infiltrated by CELF2-deficient GB cells confirmed this membrane localization, and revealed strong AHNAK expression in tumor cells juxtaposed to CD31+ endothelial structures (Fig. 6D), suggesting close physical interactions between tumor and vascular components in the assembloid microenvironment. To determine whether AHNAK is functionally required for the endothelial tropism of CELF2-deficient GB cells, we performed stably knockdown of AHNAK using three independent AHNAK-targeting shRNA sequences. Western blot analysis confirmed efficient AHNAK knockdown in all clones, with a near-complete depletion in clones B and C, which were selected for CO+EO co-culture experiments (Fig. 6E). CELF2-deficient cells transfected with a non-targeting shRNA were used as controls (Fig. 6E).

Following three weeks of co-culture, CO+EO assembloids were harvested, sectioned, and analyzed. The GFP signal from tumor cells enabled the visualization and spatial mapping of their distribution within the CO+EO assembloid (Fig. 6F). As expected, CELF2-deficient cells expressing control shRNA displayed pronounced endothelial tropism, preferentially infiltrating CD31^+^ vascular regions (Fig. 6F, upper line). In contrast, AHNAK-depleted cells formed peripheral aggregates and failed to efficiently infiltrate or distribute within the endothelial compartment, resulting in markedly reduced tumor occupancy in these regions (Fig. 6F, G). These findings demonstrate that AHNAK is a critical effector of endothelial tropism in CELF2-deficient GSCs. Moreover, they reveal that silencing AHNAK was sufficient to disrupt their vascular affinity and partially reprogram GB cell tropism within the CO+EO environment. More broadly, these results underscore the relevance of AHNAK as a functional mediator of GB cell–vascular interactions and highlight the utility of CO+EO assembloids as a platform for dissecting the molecular mechanisms governing GB tropism within complex cortico-endothelial environments.

**Figure 6.**
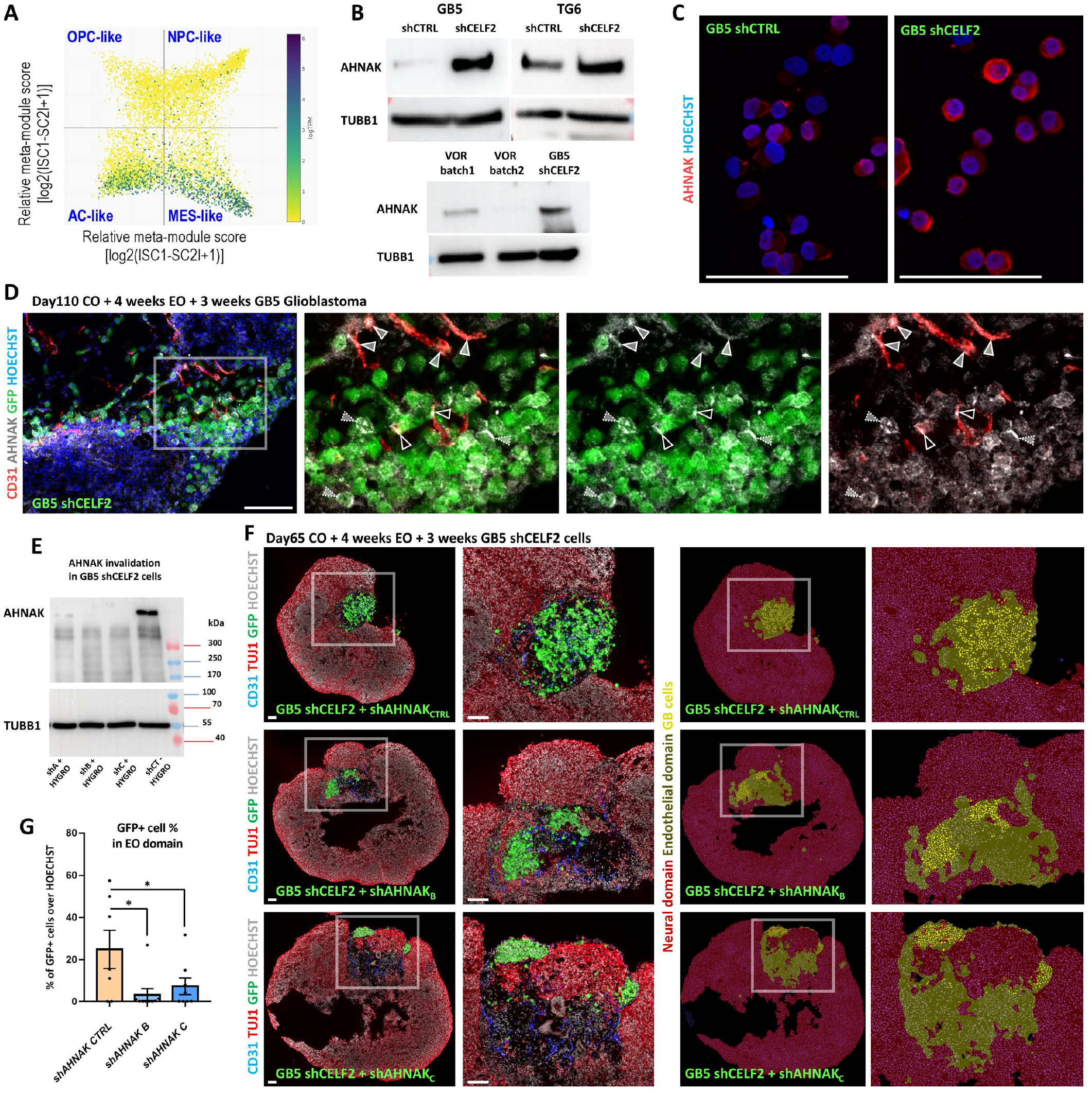
AHNAK-dependent tropism and distribution of CELF2-deprived GB cells in CO+EOs. **A)** Two-dimensional representation of AHNAK expression in distinct glioblastoma cellular states. Each quadrant corresponds to one cellular state and the blue dots represent to the highest AHNAK expression. This representation was established using the single cell portal of the Broad Institute (https://singlecell.broadinstitute.org). **B)** Western blot quantification of AHNAK protein levels in CELF2-deprived TG6 and GB5 cells, showing marked upregulation upon CELF2 loss and in endothelial cells (using EOs). **C)** Immunostaining for AHNAK (red) in GB5 shCTRL and shCELF2 cells, highlighting its membrane localization. **D)** Confocal imaging of triple immunostaining for AHNAK (white), CD31+ endothelial cells (red), and GB cells (green) in CO+EOs, showing AHNAK expression in both endothelial cells and GB cells. **E)** Western blot validation of AHNAK downregulation in three distinct clones of GB5 shCELF2 cells using specific shRNAs. **F**,**G)** Triple immunostaining for CD31 (blue; endothelial cells), TUJ1 (red; neuronal cells), and GFP (green; GB cells) in CO+EOs co-cultured with GB5 shCELF2 cells expressing a control scramble shRNA (shAHNAK_CTRL_) or AHNAK-targeting shRNAs (shAHNAK_B_ or shAHNAK_C_), with representative images of automated detection of endothelial domains and GFP+ cells on the right. Compared to control shCELF2 cells, AHNAK-depleted shCELF2 cells tend to cluster at the periphery of the endothelial domain instead of spreading throughout. Graph in G shows the percentage of GFP+ GB cells inside endothelial domains under the different conditions. Scale bars: 100 µm.

## DISCUSSION

### Modeling the glioblastoma microenvironment using a human cortico-endothelial assembloid system

GB aggressiveness, including therapeutic resistance, is driven by intrinsic tumor cell plasticity and by dynamic interactions with various cell types in the complex GB microenvironment. In particular, neurons and endothelial cells critically influence tumor cell behavior and contribute to disease progression by interacting with GB cells and secreting factors that maintain a GSC state [27] [28]. GB is characterized by a highly developed vasculature, the vessels are often dysfunctional, leading to hypoxic and necrotic tumor cores also supporting GSC maintenance and tumor aggressiveness [29]. Several 3D in vitro models have been developed to mimic human GB architecture [30]. Patient-derived GB organoids retain key histopathological features of the original tumors, including nuclear atypia, high mitotic activity, and pleomorphic nuclei [31], along with glial and GSC markers like SOX2 and OLIG2. However, these models generally lack healthy brain cells, particularly neurons, and only sporadically contain CD31-positive endothelial cells, thus limiting the study of tumor-host interaction at the neurovascular interface, where critical mechanisms of invasion and microenvironmental remodeling occur.

We developed a human iPSC-derived cortico-endothelial assembloid model (CO+EOs) that combines cortical and endothelial tissues in a 3D context. Although this model lacks full brain cell diversity, it offers key advantages. It provides a human-specific environment for patient-derived GB cells, avoiding xenograft limitations, and includes mature neurons and vascular-like structures, two critical cell types for GB physiopathology. Unlike HUVEC-based models, it better reflects BBB properties, including tight junction formation and low permeability [32]. More complex and functional BBB models incorporating HBMECs, astrocytes, and pericytes have been developed [33], yet their integration into 3D organoids remains technically challenging [34].

Adapting and extending previously established protocols [19, 20, 35], we developed a novel CO+EO assembloid system in which cortical and endothelial tissues co-develop and progressively mature over several weeks. This extended co-culture period supports the emergence of brain-like cytoarchitecture, including neuronal layers interspersed with endothelial networks expressing BBB markers such as CLDN5, OCLN, and ZO-1. These features are consistent with recent studies showing that similar organoid-based assembloids can recapitulate fundamental aspects of human BBB structure and function [36]. The CO+EO model thus offers a physiologically relevant platform to explore neurovascular interactions and, specifically, it enables detailed investigation of the mechanism by which GB cells infiltrate and remodel brain tissue at the tumor-brain interface, a critical step in tumor progression that remains poorly understood in traditional models.

### CELF2 modulates GB cell identity, tissue tropism, and interaction with the microenvironment

GB comprises a heterogeneous population of tumor cells endowed with varying tumorigenic potential [37-39]. In this study, we focused on GSC populations differing in their expression of the RNA-binding protein CELF2 to determine its role in modulating malignant cell behaviors. Our CO+EO model revealed that CELF2 governs glioblastoma cell tropism, with CELF2-positive cells preferentially infiltrating neural tissue, whereas CELF2-deficient cells show enhanced tropism toward endothelial structures These findings are consistent with histological observations in patient samples, where CELF2-expressing GSCs are enriched in mitotically active and poorly vascularized tumor zones [10]. Transcriptomic profiling further revealed that CELF2-expressing GB cells possess a neuronal/synaptic gene signature [17], aligning with transcriptomic profiles of invasive GB cells that integrate into neural circuits at the tumor margin [26]. In contrast, CELF2-deficient cells display a mesenchymal and pro-angiogenic gene expression profile, including upregulation of integrin signaling pathways.

Importantly, the CO+EO model revealed that only CELF2-positive GB cells disrupted tight junction protein expression (e.g., CLDN5) in regions of direct contact with endothelial cells, indicating that the ability to impair vascular integrity is restricted to specific GB subpopulations. Collectively, our findings indicate that CELF2 promotes a neural progenitor-like identity and directs glioblastoma cell localization toward neural tissue. Conversely, loss of CELF2 reshapes tumor–microenvironment interactions that enhances endothelial tropism while promoting a mesenchymal shift. In this context, modulating CELF2 expression influences both glioblastoma cell phenotype and its interaction with the surrounding microenvironment.

To further dissect the molecular underpinnings of this endothelial tropism, we identified the scaffold protein AHNAK as a key effector of the endothelial tropism, selectively enriched in CELF2-deficient GB cells. Confocal imaging of CO+EO assembloids revealed co-localization of AHNAK with CD31+ endothelial structures, indicating close spatial proximity and potential physical interactions between CELF2-deficient GB cells and endothelial cells. Functional silencing of AHNAK in CELF2-deficient cells partially reduced their capacity to infiltrate endothelial regions, confirming its essential role in mediating vascular affinity. Notably, AHNAK is known to associate with tight junction proteins such as CLDN5, ZO-1, and OCLN at the BBB [40], consistent with a role in facilitating endothelial interactions without directly disrupting BBB integrity. Furthermore, high AHNAK expression levels are consistent with its known barrier-stabilizing properties and it has been linked to reduced tumor growth and better clinical outcomes, while its downregulation correlates with poor prognosis [40], reinforcing its association with a less aggressive tumor phenotype.

## General Conclusion

We present a human iPSC-derived cortico-endothelial assembloid model that recapitulates GB intra-tumoral heterogeneity at the neuronal–vascular interface. This system reveals CELF2 as a key regulator of GB stem cell tropism, driving either neuroinvasive, aggressive behavior or vascular-associated, less aggressive phenotypes. CELF2-deficient cells adopt a mesenchymal identity and rely on AHNAK for vascular interaction. Our CO+EO assembloid model provides a physiologically relevant platform to study GB cell plasticity, invasion, and microenvironmental crosstalk, enabling discovery of therapeutic targets that disrupt tissue-specific tumor infiltration.

## Supporting information

neural and mesenchymal signatures

## Ethical statements

The studies were conducted in accordance with the recognized ethical standards of our institutions (Nice University Hospital and INSERM) and were approved by our institutional review board (INSERM scientific commission and ethics committee). We obtained written informed consent from all patients for the use of their tumor for research purposes.

## Data Availability statement

The data that support the findings of this study are available from the corresponding author upon request.

## Acknowledgements

The authors thank ADeRTU and François Fauchon for their interest and their support in our research. The authors thank Arnaud Borderie, Sandrine Destrée, Marianne Goracci (Nice CHU) for IHC of human GB samples. Histological and microscopy analysis were performed in the Prism facility, “Plateforme PRISM-IBV-CNRS UMR 7277-INSERM U1091-Université Côte d’Azur”. The help of Baptiste Monterosso, Sameh Ben-Aicha and Samah Rekima is acknowledged. iPSCs culture and organoids have been produced with the help of PETRA platform.

## Author contribution

MB design of experiments, data research, contributed to discussion, wrote and edited the manuscript.

AL design of experiments, data research, contributed to discussion, wrote and edited the manuscript.

AS data research, contributed to discussion

GM data research, CP data research

LT data research, contributed to discussion,

BP data research

HC provides

TG6, edited the manuscript

EE provides TG6, edited the manuscript

MPJ provides TG6, edited the manuscript

FA provides GB human surgical resections, edited the manuscript

FBV provides GB human tissues, edited the manuscript

MS design of experiments, contributed to discussion, wrote and edited the manuscript

TV design of experiments, contributed to discussion, wrote and edited the manuscript

All authors have read and agreed to the published version of the manuscript.

## References

1. Tan, A.C., et al., Management of glioblastoma: State of the art and future directions. CA Cancer J Clin, 2020. 70(4): p. 299–312.

2. Stupp, R., et al., Radiotherapy plus concomitant and adjuvant temozolomide for glioblastoma. N Engl J Med, 2005. 352(10): p. 987–96.

3. Ghosh, N., D. Chatterjee, and A. Datta, Tumor heterogeneity and resistance in glioblastoma: the role of stem cells. Apoptosis, 2025.

4. Tang, J., M.A. Amin, and J.L. Campian, Glioblastoma Stem Cells at the Nexus of Tumor Heterogeneity, Immune Evasion, and Therapeutic Resistance. Cells, 2025. 14(8).

5. Turchi, L., et al., Tumorigenic Potential of miR-18A* in Glioma Initiating Cells Requires NOTCH-1 Signaling. Stem Cells, 2013. Jul;31(7): p. 1252–65.

6. Almairac, F., et al., ERK-Mediated Loss of miR-199a-3p and Induction of EGR1 Act as a “Toggle Switch” of GBM Cell Dedifferentiation into NANOG- and OCT4-Positive Cells. Cancer Res, 2020. 80(16): p. 3236–3250.

7. Vartanian, A., et al., GBM’s multifaceted landscape: highlighting regional and microenvironmental heterogeneity. Neuro Oncol, 2014. 16(9): p. 1167–75.

8. Markwell, S.M., et al., Necrotic reshaping of the glioma microenvironment drives disease progression. Acta Neuropathol, 2022. 143(3): p. 291–310.

9. Debruyne, D.N., et al., DOCK4 promotes loss of proliferation in glioblastoma progenitor cells through nuclear beta-catenin accumulation and subsequent miR-302-367 cluster expression. Oncogene, 2018. 37(2): p. 241–254.

10. Turchi, L., et al., CELF2 Sustains a Proliferating/OLIG2+ Glioblastoma Cell Phenotype via the Epigenetic Repression of SOX3. Cancers (Basel), 2023. 15(20).

11. Alghamri, M.S., et al., Targeting Neuroinflammation in Brain Cancer: Uncovering Mechanisms, Pharmacological Targets, and Neuropharmaceutical Developments. Front Pharmacol, 2021. 12: p. 680021.

12. Zarco, N., et al., Overlapping migratory mechanisms between neural progenitor cells and brain tumor stem cells. Cell Mol Life Sci, 2019. 76(18): p. 3553–3570.

13. Watkins, S., et al., Disruption of astrocyte-vascular coupling and the blood-brain barrier by invading glioma cells. Nat Commun, 2014. 5: p. 4196.

14. Varn, F.S., et al., Glioma progression is shaped by genetic evolution and microenvironment interactions. Cell, 2022. 185(12): p. 2184-2199.e16.

15. Westerlund, L.H., et al., Deciphering the Dialogue between Brain Tumors, Neurons, and Astrocytes. Am J Pathol, 2025.

16. Barreau, C., et al., Mammalian CELF/Bruno-like RNA-binding proteins: molecular characteristics and biological functions. Biochimie, 2006. 88(5): p. 515–25.

17. Neftel, C., et al., An Integrative Model of Cellular States, Plasticity, and Genetics for Glioblastoma. Cell, 2019. 178(4): p. 835-849.e21.

18. Zhang, S., Z. Cai, and H. Li, AHNAKs roles in physiology and malignant tumors. Front Oncol, 2023. 13: p. 1258951.

19. Wimmer, R.A., et al., Generation of blood vessel organoids from human pluripotent stem cells. Nat Protoc, 2019. 14(11): p. 3082–3100.

20. Bertacchi, M., et al., FGF8-mediated gene regulation affects regional identity in human cerebral organoids. Elife, 2024. 13.

21. Yu, W., et al., STAT3-controlled CHI3L1/SPP1 positive feedback loop demonstrates the spatial heterogeneity and immune characteristics of glioblastoma. Dev Cell, 2025.

22. Ke, C., et al., Dual antivascular function of human fibulin-3 variant, a potential new drug discovery strategy for glioblastoma. Cancer Sci, 2020. 111(3): p. 940–950.

23. Hu, Y., et al., Cell context-dependent dual effects of EFEMP1 stabilizes subpopulation equilibrium in responding to changes of in vivo growth environment. Oncotarget, 2015. 6(31): p. 30762–72.

24. Jung, J., et al., Nicotinamide metabolism regulates glioblastoma stem cell maintenance. JCI Insight, 2017. 2(10).

25. Jia, D., et al., Mining TCGA database for genes of prognostic value in glioblastoma microenvironment. Aging (Albany NY), 2018. 10(4): p. 592–605.

26. Ren, Y., et al., Spatial transcriptomics reveals niche-specific enrichment and vulnerabilities of radial glial stem-like cells in malignant gliomas. Nat Commun, 2023. 14(1): p. 1028.

27. Maleszewska, M., et al., Decoding glioblastoma’s diversity: Are neurons part of the game? Cancer Lett, 2025. 620: p. 217666.

28. Calabrese, C., et al., A perivascular niche for brain tumor stem cells. Cancer Cell, 2007. 11(1): p. 69–82.

29. Heddleston, J.M., et al., The hypoxic microenvironment maintains glioblastoma stem cells and promotes reprogramming towards a cancer stem cell phenotype. Cell Cycle, 2009. 8(20): p. 3274–84.

30. Gómez-Oliva, R., et al., Evolution of Experimental Models in the Study of Glioblastoma: Toward Finding Efficient Treatments. Front Oncol, 2020. 10: p. 614295.

31. Jacob, F., et al., A Patient-Derived Glioblastoma Organoid Model and Biobank Recapitulates Inter- and Intra-tumoral Heterogeneity. Cell, 2020. 180(1): p. 188-204.e22.

32. Fedele, G., et al., The presence of BBB hastens neuronal differentiation of cerebral organoids - The potential role of endothelial derived BDNF. Biochem Biophys Res Commun, 2022. 626: p. 30–37.

33. Bergmann, S., et al., Blood-brain-barrier organoids for investigating the permeability of CNS therapeutics. Nat Protoc, 2018. 13(12): p. 2827–2843.

34. Bergmann, N., et al., The Intratumoral Heterogeneity Reflects the Intertumoral Subtypes of Glioblastoma Multiforme: A Regional Immunohistochemistry Analysis. Front Oncol, 2020. 10: p. 494.

35. Sun, X.Y., et al., Generation of vascularized brain organoids to study neurovascular interactions. Elife, 2022. 11.

36. Dao, L., et al., Modeling blood-brain barrier formation and cerebral cavernous malformations in human PSC-derived organoids. Cell Stem Cell, 2024. 31(6): p. 818-833.e11.

37. Sottoriva, A., et al., Intratumor heterogeneity in human glioblastoma reflects cancer evolutionary dynamics. Proceedings of the National Academy of Sciences of the United States of America, 2013. 110(10): p. 4009–14.

38. Suvà, M.L. and I. Tirosh, The Glioma Stem Cell Model in the Era of Single-Cell Genomics. Cancer Cell, 2020. 37(5): p. 630–636.

39. Patel, A.P., et al., Single-cell RNA-seq highlights intratumoral heterogeneity in primary glioblastoma. Science, 2014. 344(6190): p. 1396–401.

40. Gentil, B.J., et al., Specific AHNAK expression in brain endothelial cells with barrier properties. J Cell Physiol, 2005. 203(2): p. 362–71.

