## supplemental figure and legends for "Human cortico-vascular assembloids reveal a CELF2-AHNAK-dependent switch from neuronal to endothelial tropism in glioblastoma cells"

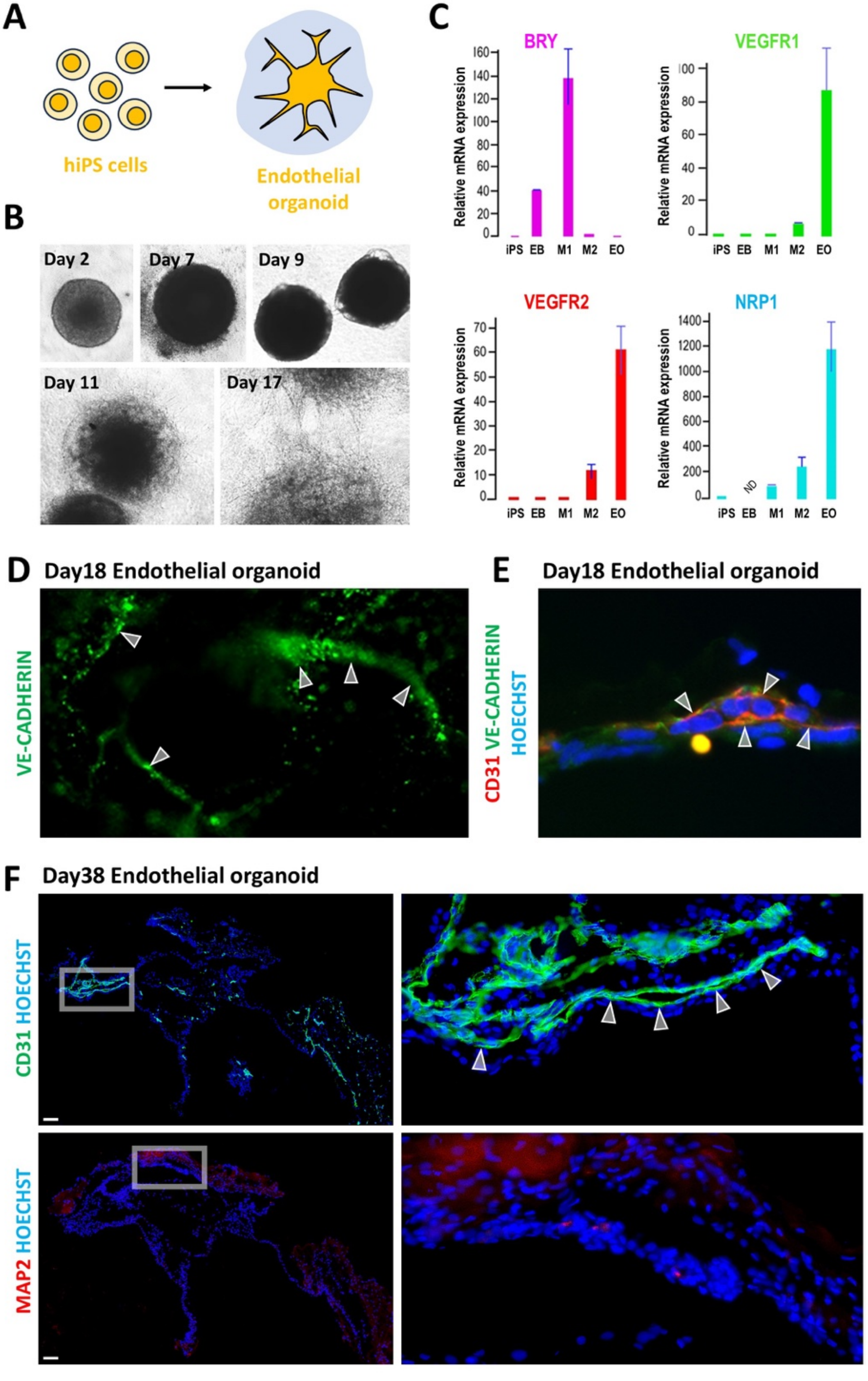

A
hiPS cells
Endothelial organoid
B
Day 2
Day 7
Day 9
Day 11
Day 17
C
BRY
VEGFR1
VEGFR2
NRP1
Relative mRNA expression
iPS EB M1 M2 EO
ND
D
Day18 Endothelial organoid
VE-CADHERIN
E
Day18 Endothelial organoid
CD31 VE-CADHERIN HOECHST
F
Day38 Endothelial organoid
CD31 HOECHST
MAP2 HOECHST

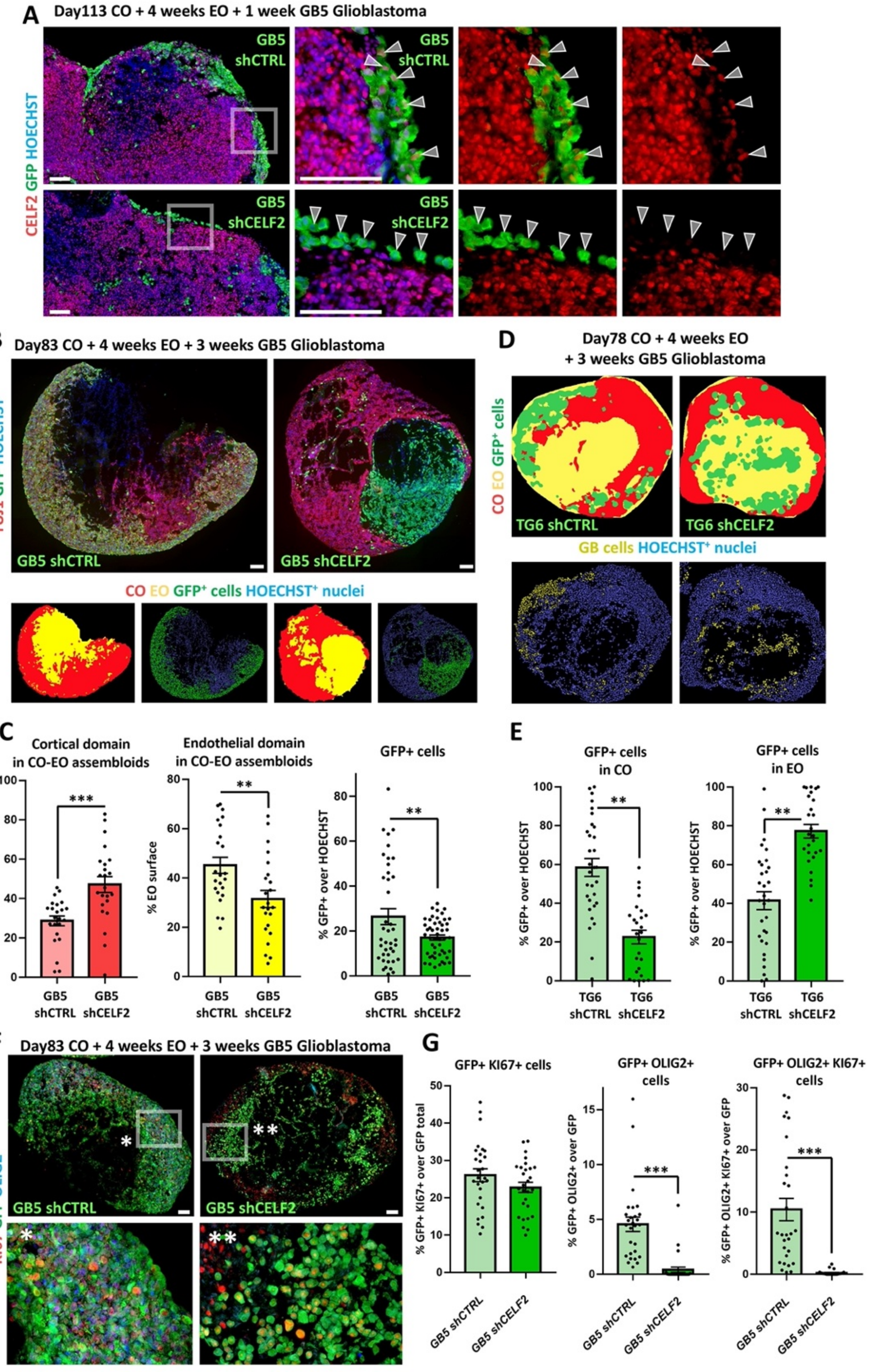

A
Day113 CO + 4 weeks EO + 1 week GB5 Glioblastoma
CELF2 GFP HOECHST
GB5 shCTRL
GB5 shCELF2
B
Day83 CO + 4 weeks EO + 3 weeks GB5 Glioblastoma
TUJ1 GFP HOECHST
GB5 shCTRL
GB5 shCELF2
CO EO GFP+ cells HOECHST+ nuclei
D
Day78 CO + 4 weeks EO + 3 weeks GB5 Glioblastoma
CO EO GFP+ cells
TG6 shCTRL
TG6 shCELF2
GB cells HOECHST+ nuclei
C
Cortical domain in CO-EO assembloids
% MAP2+ surface
Endothelial domain in CO-EO assembloids
% EO surface
GFP+ cells
% GFP+ over HOECHST
GB5 shCTRL
GB5 shCELF2
E
GFP+ cells in CO
GFP+ cells in EO
% GFP+ over HOECHST
TG6 shCTRL
TG6 shCELF2
F
Day83 CO + 4 weeks EO + 3 weeks GB5 Glioblastoma
KI67 GFP OLIG2
GB5 shCTRL
GB5 shCELF2
G
GFP+ KI67+ cells
% GFP+ KI67+ over GFP total
GFP+ OLIG2+ cells
% GFP+ OLIG2+ over GFP
GFP+ OLIG2+ KI67+ cells
% GFP+ OLIG2+ KI67+ over GFP
GB5 shCTRL
GB5 shCELF2

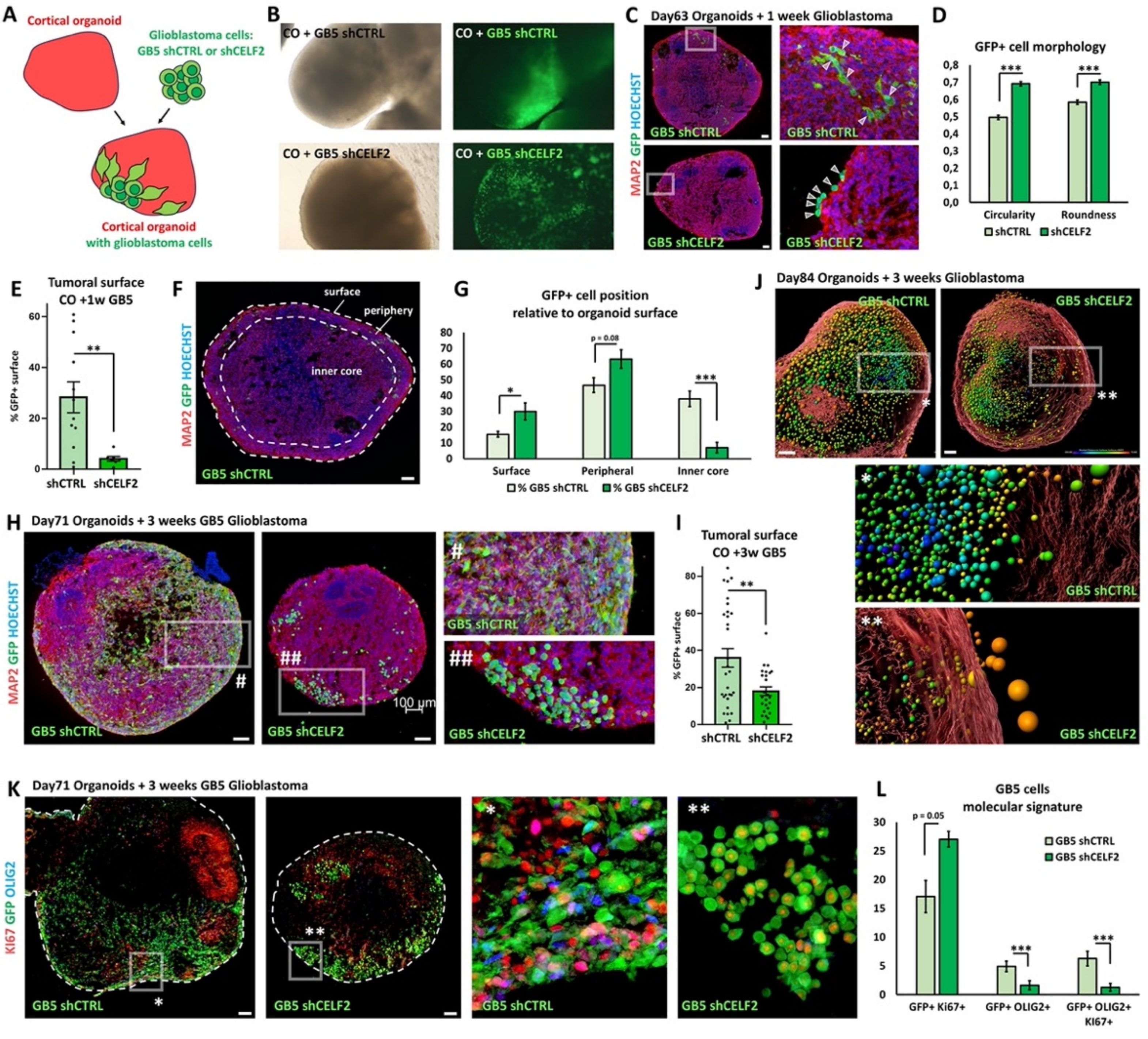
A
Cortical organoid
Glioblastoma cells:
GB5 shCTRL or shCELF2
Cortical organoid
with glioblastoma cells
B
CO + GB5 shCTRL
CO + GB5 shCTRL
CO + GB5 shCELF2
CO + GB5 shCELF2
C
Day63 Organoids + 1 week Glioblastoma
MAP2 GFP HOECHST
GB5 shCTRL
GB5 shCTRL
GB5 shCELF2
GB5 shCELF2
D
GFP+ cell morphology
0,8
0,7
0,6
0,5
0,4
0,3
0,2
0,1
0,0
***
***
Circularity
Roundness
shCTRL
shCELF2
E
Tumoral surface
CO +1w GB5
% GFP+ surface
60
40
20
0
**
shCTRL
shCELF2
F
MAP2 GFP HOECHST
surface
periphery
inner core
GB5 shCTRL
G
GFP+ cell position
relative to organoid surface
80
70
60
50
40
30
20
10
0
*
p = 0.08
***
Surface
Peripheral
Inner core
% GB5 shCTRL
% GB5 shCELF2
H
Day71 Organoids + 3 weeks GB5 Glioblastoma
MAP2 GFP HOECHST
#
GB5 shCTRL
##
100 µm
GB5 shCELF2
#
GB5 shCTRL
##
GB5 shCELF2
I
Tumoral surface
CO +3w GB5
% GFP+ surface
80
60
40
20
0
**
shCTRL
shCELF2
J
Day84 Organoids + 3 weeks Glioblastoma
GB5 shCTRL
GB5 shCELF2
*
**
*
GB5 shCTRL
**
GB5 shCELF2
K
Day71 Organoids + 3 weeks GB5 Glioblastoma
KI67 GFP OLIG2
GB5 shCTRL
*
**
GB5 shCELF2
*
GB5 shCTRL
**
GB5 shCELF2
L
GB5 cells
molecular signature
30
25
20
15
10
5
0
p = 0.05
***
***
GB5 shCTRL
GB5 shCELF2
GFP+ Ki67+
GFP+ OLIG2+
GFP+ OLIG2+
KI67+

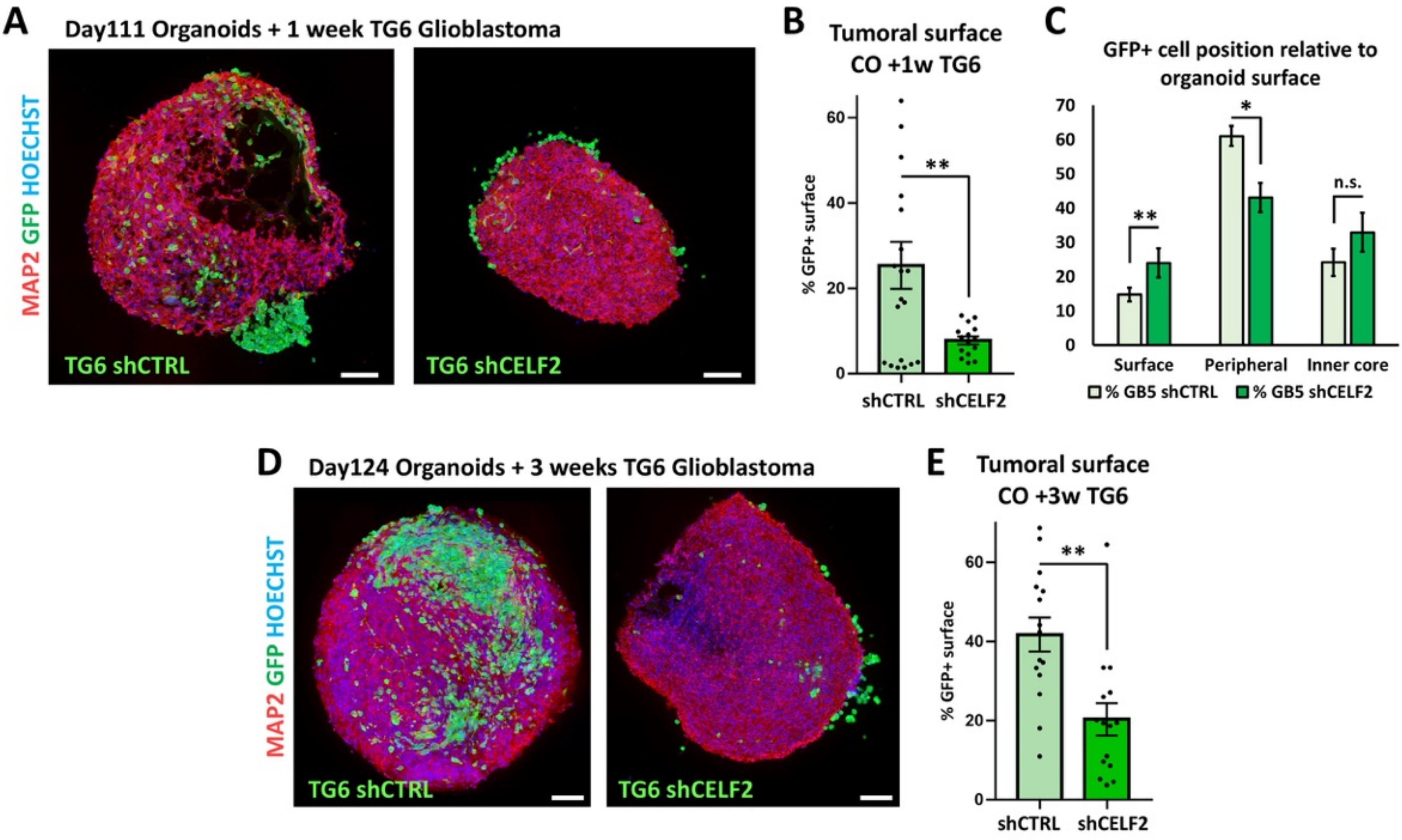
A
Day111 Organoids + 1 week TG6 Glioblastoma
MAP2 GFP HOECHST
TG6 shCTRL
TG6 shCELF2
B
Tumoral surface
CO +1w TG6
% GFP+ surface
0
20
40
60
**
shCTRL
shCELF2
C
GFP+ cell position relative to organoid surface
0
10
20
30
40
50
60
70
**
*
n.s.
Surface
Peripheral
Inner core
% GB5 shCTRL
% GB5 shCELF2
D
Day124 Organoids + 3 weeks TG6 Glioblastoma
MAP2 GFP HOECHST
TG6 shCTRL
TG6 shCELF2
E
Tumoral surface
CO +3w TG6
% GFP+ surface
0
20
40
60
**
shCTRL
shCELF2

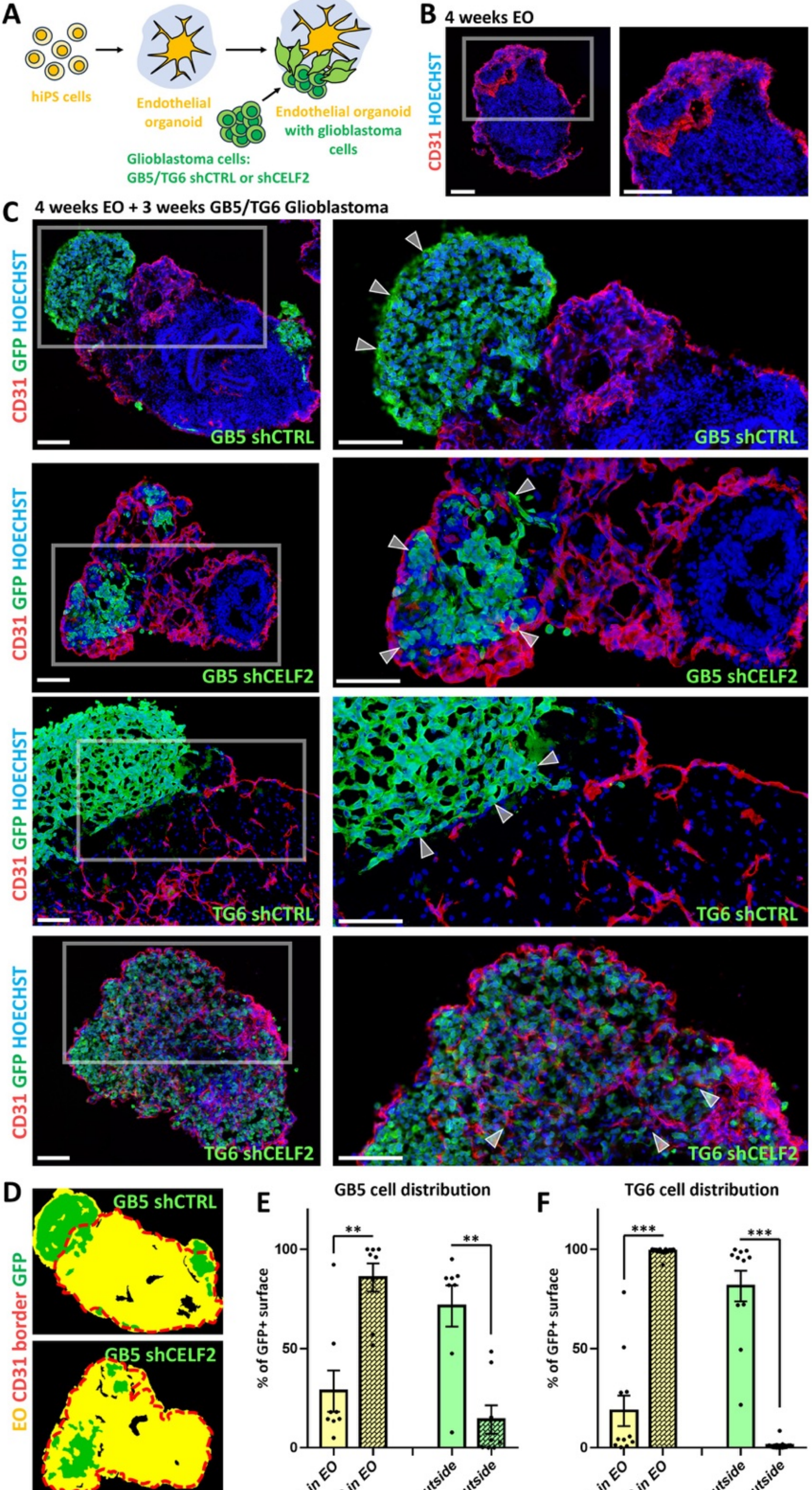
A
hiPS cells
Endothelial organoid
Endothelial organoid with glioblastoma cells
Glioblastoma cells: GB5/TG6 shCTRL or shCELF2
B 4 weeks EO
CD31 HOECHST
C 4 weeks EO + 3 weeks GB5/TG6 Glioblastoma
CD31 GFP HOECHST
GB5 shCTRL
GB5 shCELF2
TG6 shCTRL
TG6 shCELF2
D
EO CD31 border GFP
E GB5 cell distribution
F TG6 cell distribution
% of GFP+ surface
0
50
100
**
***
shCTRL in EO
shCELF2 in EO
shCTRL outside
shCELF2 outside

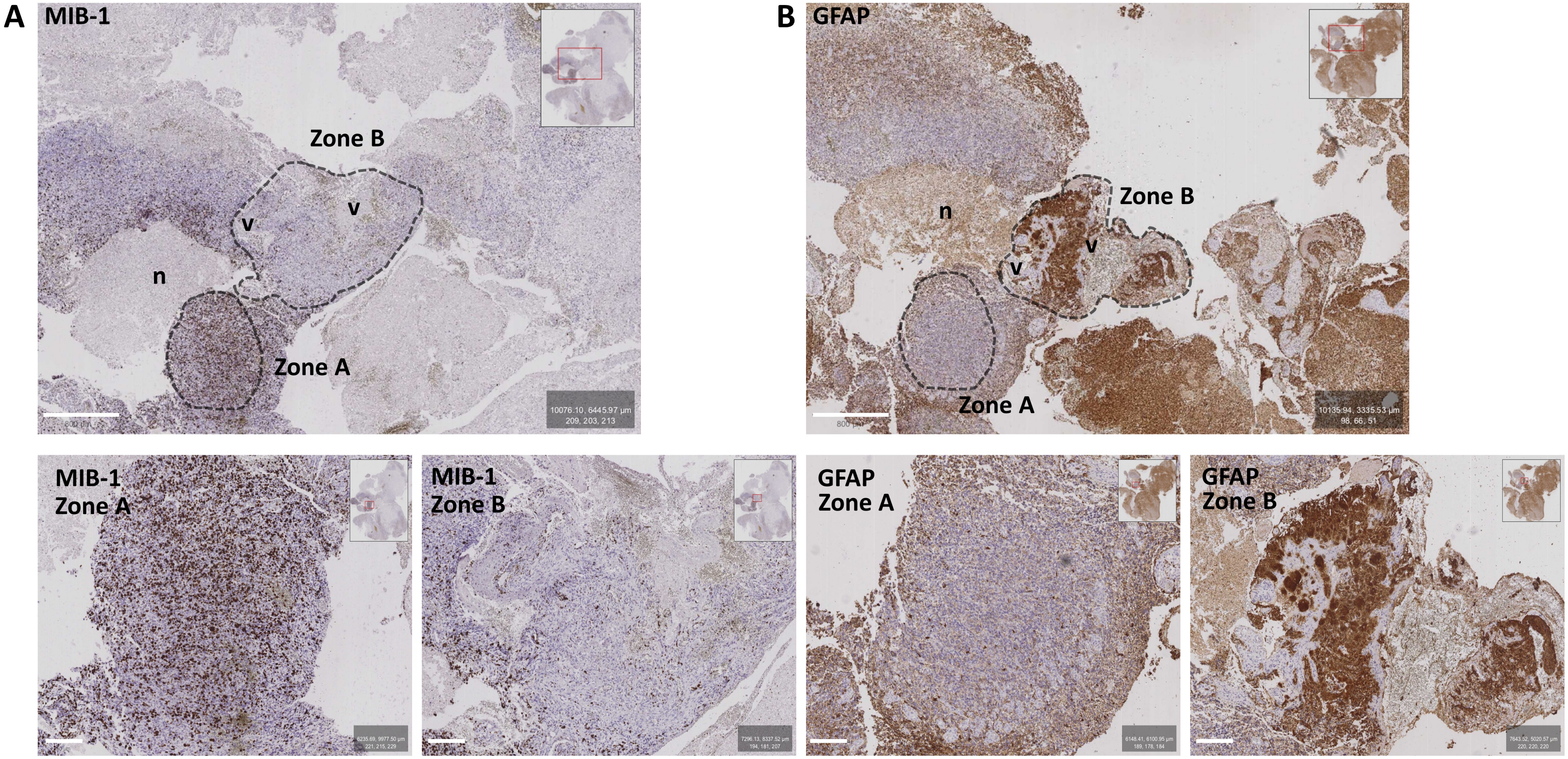
A
MIB-1
Zone B
v
v
n
Zone A
B
GFAP
Zone B
n
v
v
Zone A
MIB-1
Zone A
MIB-1
Zone B
GFAP
Zone A
GFAP
Zone B

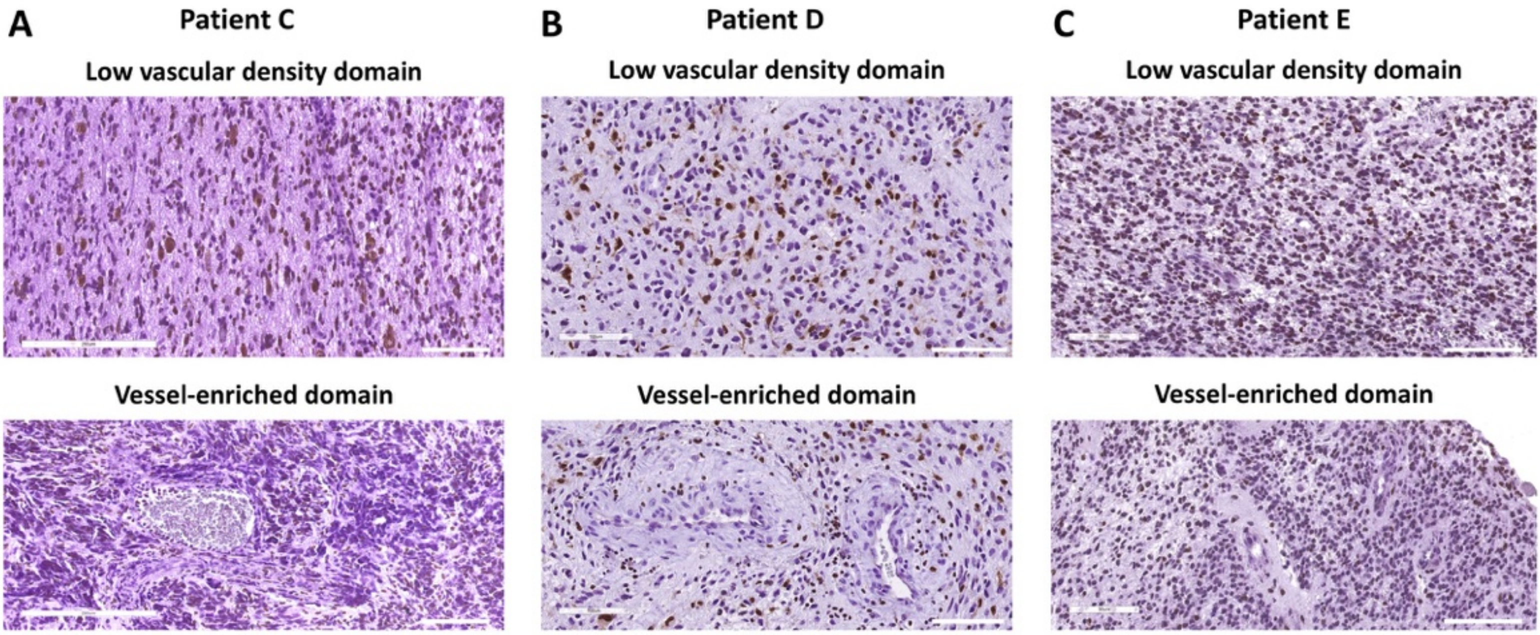
A
Patient C
Low vascular density domain
Vessel-enriched domain
B
Patient D
Low vascular density domain
Vessel-enriched domain
C
Patient E
Low vascular density domain
Vessel-enriched domain

### Supplemental figure legends

**Suppl. Figure 1. Efficient *in vitro* induction of mesodermal tissue and endothelial organoids. A–B)** Endothelial organoids (EOs) were derived from hiPSCs (A) *via* embryoid body formation and subsequent culture in VEGF/FGF2-containing medium, resulting in the emergence of elongated, multicellular structures reminiscent of blood vessels (B). **C)** Real-time qRT-PCR analysis of mesodermal and endothelial markers showing robust mesodermal induction (BRACHYURY, BRY) followed by endothelial maturation, as indicated by upregulation of VEGFR1, VEGFR2, and NRP1. **D–F)** Immunostaining of *in vitro*–differentiated endothelial cells in EOs showing expression of VE-CADHERIN (green, D and E) and CD31 (red in E; green in F), with no detectable expression of the neuronal marker MAP2 (red in F). Scale bars: 100 µm.

**Suppl. Figure 2. CELF2-dependent distribution and stemness identity of GB cells in CO+EOs. A)** Immunostaining for CELF2 (red) and GFP (green) in shCTRL (upper row) and shCELF2 (lower row) GB5 cells one week after co-culture with CO+EOs. **B,C)** TUJ1 (red) and GFP (green) immunostaining of CO+EOs and automated recognition of tumor cells (green), cortical domains (red), or endothelial domains (yellow); graphs in C display the percentages of each surface type in human assembloids 3 weeks after GB invasion: MAP2+ cortical surface (left), endothelial surface (middle), and percentage of GFP+ GB cells (right graph). **D,E)** Automated recognition of tumor (green), endothelial (yellow), and cortical (red) surfaces in CO+EOs 3 weeks after invasion with TG6 shCTRL or TG6 shCELF2 cells, showing that these different cell lines exhibit the same tropism toward mesodermal tissue upon CELF2 downregulation. Graphs in E show the percentage of GFP+ cells found in the CO or EO domains. **F,G)** Triple immunostaining for GFP (green), KI67 (red), and OLIG2 (blue) in shCTRL and shCELF2 assembloids after 3 weeks of co-culture (F). Graphs in G show the percentages of GFP+ cells positive for KI67, OLIG2, or both KI67 and OLIG2, indicating marked loss of OLIG2+ GSC identity upon CELF2 downregulation. Scale bars: 100 µm.

**Suppl. Figure 3. Glioblastoma-invaded COs reveal CELF2-dependent dynamics of GB cell morphology, migration, proliferation, and stemness. A)** Schematic representation of cortico-glioblastoma co-culture integrating GB5 control (shCTRL) or CELF2-deprived (shCELF2) GB5 cells into COs. **B)** Brightfield images showing COs one week after co-culture with shCTRL GB5 cells (top row) or with shCELF2 GB5 cells (lower row); CELF2-deprived cells can be seen on the organoid surface and with round morphology. **C-E)** MAP2 (red) and GFP (green) immunostaining of day 63 COs supplemented with shCTRL and shCELF2 GB5 cells after one week of co-culture. White arrowheads in high-magnification insets point to GFP+ GB5 cells; note that shCELF2 cells exhibit a round morphology and are primarily distributed on the external surface of the COs. ImageJ quantification of cellular morphology (circularity and roundness) is shown in graph D, while graph E shows quantification of the GFP+ surface area in organoid sections after one week of co-culture. **F,G)** Evaluation of GB cell position in organoid sections, classified as on the surface, in the periphery (100 µm from the surface), or in the inner core. The gr[illegible]
shCELF2 assembloids after one week of co-culture. **H,I)** MAP2 (red) and GFP (green) immunostaining of assembloids after three weeks of co-culture. The graph in I shows quantification of the GFP+ surface area. **J)** 3D reconstruction of tissue-cleared COs invaded by shCTRL or shCELF2 GB5 cells, as indicated; automated cell recognition in Imaris allowed measurement of the distance of cells (dots) from the organoid surface; blue to red color code indicates far to close to the surface, respectively. **K,L)** Triple immunostaining for GFP (green), KI67 (red), and OLIG2 (blue) in shCTRL and shCELF2-supplemented COs after three weeks. The percentages of GFP+ cells positive for KI67, OLIG2, or both KI67 and OLIG2 are shown in graph L. Scale bars: 100 µm.

**Suppl. Figure 4. TG6 GB cells show impaired invasiveness and migratory potential in COs. A-C)** MAP2 (red) and GFP (green) immunostaining of day 111 cortical organoids (COs) supplemented with shCTRL (left) and shCELF2 (right) TG6 cells after one week of co-culture. Quantification of GFP+ tumoral surface area after one week of co-culture is shown in B, while graph C displays the position of GB ce[illegible]
peripheral outer shell, or in the inner core) for TG6 shCTRL and shCELF2 COs after one week of co-culture. **D,E)** MAP2 (red) and GFP (green) immunostaining of COs after three weeks of co-culture with TG6 control or CELF2-deprived cells. The graph in E shows quantification of the GFP+ area in COs. Scale bars: 100 µm.

**Suppl. Figure 5. GB cell invasive potential in endothelial organoids. A)** Schematic representation of endothelial organoids (EOs; yellow) and GB cells (green) in co-culture. **B,C)** CD31 (red; endothelial cells) and GFP (green; tumoral cells) immunostaining of EOs after 4 weeks of in vitro differentiation (B) and 3 weeks of co-culture with GB cells (C). Panel C shows exemplificative organoid sections of EOs co-cultured with GB5 or TG6 shCTRL and shCELF2 cells. Note that shCELF2 GB5 and TG6 cells exhibit tropism toward endothelial tissues and demonstrate high invasive potential into CD31+ domains, whereas shCTRL cells tend to form peripheral aggregates and invade less. Arrowheads indicate clusters of GFP+ cells. **D-F)** HALO software automated recognition of endothelial domains (yellow) and GFP+ GB cells (green). Graphs in E and F show the percentages of GB5 (E) and TG6 (F) GFP+ cells found inside or outside the borders of endothelial areas (red lines in D). Scale bars: 100 µm.

**Suppl. Figure 6. Varying degrees of vascular enrichment in mitotic *versus* non-mitotic zones in GB patient samples. A,B)** Immunohistochemical staining of GB samples showing the mitotic marker MIB-1 (brown in panel A) and the astrocyte differentiation marker GFAP (brown in panel B), overlaid on hematoxylin-eosin (H&E) staining, which highlights nuclei in violet. The upper low-magnification images illustrate intra-tumoral heterogeneity, with mitotic regions (Zone A) displaying sparse vasculature, and non-mitotic regions (Zone B) showing abnormal, dense vasculature (v) adjacent to necrotic areas (n). High-magnification images reveal strong MIB-1 labeling in mitotic zones and high GFAP labeling in non-mitotic zones. Scale bars: 800 µm (low magnification) and 200 µm (high magnification).

**Suppl. Figure 7. CELF2+ cell distribution in GB regions with varying vascular enrichment. A-C)** Immunohistochemistry staining of GB samples from three patients, showing CELF2 (brown) over hematoxylin-eosin staining that highlights nuclei (violet). The upper row displays areas with few and small blood vessels (low vascular density domains), while the lower row illustrates highly vascularized areas with large blood vessels (vessel-enriched domains). Quantification of CELF2+ cells in different domains is shown in Figure 3. Scale bars: 100 µm.
