## supplemental table for "Human cortico-vascular assembloids reveal a CELF2-AHNAK-dependent switch from neuronal to endothelial tropism in glioblastoma cells"

Supplemental table 1. Antibodies used in immunostaining experiments in this study

| Primary antibody | species | Concentration used | Antigen retrieval | Brand | ref |
|---|---|---|---|---|---|
| CD31 | mouse | 1/1000 | NO | Abcam | Ab9498 |
| VE-Cadherin | Rabbit | 1/2000 | NO | GENTEX | GTX132982 |
| CELF2 | Rabbit | 1/1000 | YES | MERCK | HPA035813 |
| Cleaved Caspase 3 | Rabbit | 1/1000 | YES | Cell signaling | #9661 |
| CLDN5 | Rabbit | 1/1000 | NO | Abcam | AB 131259 |
| OCLN | rabbit | 1/1000 | NO | GENTEX | 114949 |
| ZO-1 | rabbit | 1/500 | NO | Gentex | 636399 |
| AHNAK | RABBIT | 1/500 ICC; 1/200 WB | NO | INVITROGEN | PA5-53890 |
| SERPING1 | RABBIT | 1/1000 | NO | INVITROGEN | MA5-46954 |
| CHI3L1 | Rabbit | 1/1000 | NO | ABCAM | Ab77528 |
| DCX | Rabbit | 1/1000 | NO | Cell signaling technology | 40619 |
| TUBBI | mouse | 1/10 000 (WB) | NO | SIGMA | T7816 |
| GFP | Chicken | 1/1000 | NO | Abcam | Ab13970 |
| KI67 | Rabbit | 1/1000 | YES | Abcam | Ab15580 |
| MAP2 | Mouse | 1/1000 | NO | Sigma | M4403 |
| OLIG2 | Mouse | 1/1000 | YES | Millipore | MABN50 |
| TUJ1 | Rabbit | 1/1000 | NO | BioLegend | 802001 |

Supplemental table 2. Sequences for the primers used in this study

| Gene | Forward primer | Reverse primer | EFFICIENCY |
|---|---|---|---|
| AHNAK | 5'-CCTGAAAGGGCCTCGGATTT-3' | 5'- GCCTTGAACACCAATGCCTG-3' | 101% |
| BRY | 5'-ATGATGGAGGAACCCGGAGA-3' | 5'-TGGTTCCAGGAAGAAGCCAC -3' | 99% |
| VE-Cadherin | 5'- ATGCGGCTAGGCATAGCATT | 5'- TGTGACTCGGAAGAACTGGC-3' | 97% |
| NRP1 | 5'- GGGGCTCTCACAAGACCTTC -3' | 5'- GATCCTGAATGGGTCCCGTC -3' | 100% |
| VEGFR1 | 5'- acaaggcaagaaaccaagac-3' | 5'- cctcctcttcctcaacatcac-3' | 100% |
| VEGFR2 | 5'- tcagcaggatggcaaagac-3' | 5'- catacttcctcctcctccatac-3' | 98% |
| 36B4 | 5'-CTACAACCCTGAAGAAGTGCTTG-3' | 5'-CAATCTGCAGACAGACACTGG-3' | 102% |
