## supplemental materials and methods for "Human cortico-vascular assembloids reveal a CELF2-AHNAK-dependent switch from neuronal to endothelial tropism in glioblastoma cells"

### SUPP MATERIALS AND METHODS

#### *hiPSC culture*

Human induced pluripotent stem cells (hiPSCs) used in this study are the HMGU1 cell line, derived from fibroblasts established from normal foreskin of a neonatal male (ATCC number CRL-2522, designation BJ), mycoplasma-free and kindly provided by Dr. Drukker. Material Transfer Agreement (MTA) approval was obtained from the Helmholtz Zentrum München (HMGU), Germany. HMGU1 hiPSCs were cultured on Matrigel-coated plates (Corning, 354234; 5–10 µL Matrigel dissolved in 1 mL cold DMEM/F12 per well of a six-well culture plate) in mTeSR1 medium (STEMCELL Technologies, #85850). Medium was changed daily. HMGU1 cells were passaged with Versene (Thermo Fisher Scientific, 15040066) as previously described (Beers et al., 2012); briefly, cells were washed with 1 mL PBS1x (Thermo Fisher Scientific, 14190169; or Sigma, D8537), treated with Versene for up to 5 minutes, then gently detached by pipetting with 1 mL mTeSR1 medium. Alternatively, for single-cell dissociation and cell counting, HMGU1 cells were passaged using Accutase (Sigma, A6964). Single-cell passaging required the addition of 10 µM dihydrochloride ROCK inhibitor Y-27632 (MedChemExpress, HY-10583; or STEMCELL Technologies, #72304) to prevent apoptosis; ROCK inhibitor was removed 24 hours after dissociation.

#### *Glioblastoma cell line culture*

Patient-derived GB5 cells were isolated from surgical resections of primary human GBMs provided by the Department of Neurosurgery of the University Hospital of Nice.TG6 cells were kindly supplied by Hervé Chneiweiss (Université Pierre et Marie Curie, Paris). Patient-derived cell cultures were enriched for glioma stem cells and grown as spheroids as previously described [10, 19]. Briefly, spheroids were maintained in NS34+ medium supplemented with EGF and FGF-2 (Miltenyi Biotec; 130-093-841) at a final concentration of 20 ng/mL each. NS34+ medium consisted of DMEM/F12 (ThermoFisher Scientific, #31331028) supplemented with 10 mM Glutamine (ThermoFisher Scientific, 35050061), 10 mM HEPES (ThermoFisher Scientific, 15630056), 0.025% sodium bicarbonate (7.5% solution; ThermoFisher, 5080094), N-2 Supplement 100X (ThermoFisher Scientific, 17502-048), G5 Supplement (ThermoFisher Scientific, 17503012), and B-27 Supplement 50X, minus vitamin A (ThermoFisher Scientific, 12587-010). shCTRL and shCELF2 patient-derived cells were produced as previously described [10].

#### *hiPSC and Glioblastoma cell line quality controls*

HMGU1 cells and glioma stem cells were routinely tested for mycoplasma contamination once per year during the amplification phase and preparation of multiple cryovial aliquots, which were subsequently used in downstream experiments over the following months. Mycoplasma detection was performed via conventional PCR using cell culture supernatant that had been in contact with the cells for at least 48 hours. The reaction mix (50 µL) included: 5 µL of 10X PCR buffer, 1 µL of 10 mM dNTPs, 1 µL each of Myco_F1 (5'-CTTCATCGACTTTCAGA-3') and Myco_R1 (5'-ACACCATGGGAGCTGGTAAT-3') primers at 10 µM, 0.5 µL Taq polymerase, 5 µL of culture supernatant, and 36.5 µL of nuclease-free water. The expected amplicon size was approximately 600 bp.

Genetic and chromosomal integrity of HMGU1 cells was assessed using the hPSC Genetic Analysis Kit (Stem Cell Technologies; #07550) according to the manufacturer's instructions. Two additional human iPSC lines—WT-T12 (a kind gift from Dr. Magdalena Laugsch) and PGP1 (purchased from Synthego) were included as controls for comparative analysis. Briefly, hiPSC cell pellets were lysed in a buffer containing 1M Tris-HCl pH 8.0, 5M NaCl, 0.5M EDTA, and 10% SDS in Milli-Q water, supplemented with 10 mg/mL Proteinase K (Sigma, 0311580), and incubated at 58°C for 30 minutes. Genomic DNA was extracted via isopropanol precipitation followed by washing with 70% ethanol. Frequent mutations were detected in the HMGU1 cell line, and extended in vitro culture led to the accumulation of chromosomal abnormalities in vulnerable genomic regions. To minimize this, HMGU1 cells were not used beyond passage 26. Pluripotency of the hiPSC lines was confirmed by OCT4 immunostaining.

***Neural differentiation of cortical organoids (COs)***

HMGU1 cells were differentiated into neural organoids as described previously [20]. Briefly, HMGU1 cells were seeded in Matrigel-coated 24-well plates at a density of 25,000–50,000 cells per well and cultured in mTeSR1 or in mTeSR+ medium until reaching high confluency. When cells were 90–95% confluent, hiPSCs were washed with pre-warmed DMEM/F12, and the medium was switched to neural progenitor patterning medium (NPPM). NPPM consisted of DMEM/F12 supplemented with N2 supplement (ThermoFisher Scientific, 17502001, B-27 Supplement 50X minus vitamin A (ThermoFisher Scientific, 12587-010), GlutaMAX (ThermoFisher Scientific, 35050038), Non-Essential Amino Acids (NEAA; ThermoFisher Scientific, 11140-035), Sodium Pyruvate (NaPyr; ThermoFisher Scientific, 11360070), 50 µM 2-mercaptoethanol (ThermoFisher Scientific, 31350-010), 2 µg/mL heparin (Sigma, H3149), and Penicillin/Streptomycin (Biozol, ECL-EC3001D), which could alternatively be replaced with Antibiotic/Antimycotic solution (Sigma, A5955). Neural induction was enhanced by adding a BMP inhibitor (0.25 µM LDN-193189; Sigma, SML0559-5MG), a TGFβ inhibitor (5 µM SB-431542; Sigma, S4317-5MG), and a WNT inhibitor (2 µM XAV-939; StemCell Technologies, 72674). On day 7 (maximum day 8), cells were dissociated using Accutase and seeded into 96-well U-bottom plates (Corning, 7007; or Sarstedt, 83.3925500) at a concentration of 60,000 cells/well in NPPM medium supplemented with ROCK inhibitor. A gentle centrifugation of the 96-well plate (1 minute at 1,000 rpm) facilitated cell aggregation at the bottom of the wells. The following day, embryoid bodies (EBs) were collected and embedded in Matrigel droplets (Corning, 354234; 70 µL Matrigel for 16 EBs) to form "cookies," which were placed on parafilm and incubated for 30 minutes at 37 °C to allow Matrigel solidification, then transferred into SpinΩ mini-rotors (ref 67 from FGF8 paper) in NPPM medium (still supplemented with SB + LDN + XAV; 2.5 mL per well in a 12-well plate). As an alternative to SpinΩ mini-rotor, we used a $CO_2$-resistant orbital shaker (Fisher Scientific, catalog number/model: 88881102; accessories: aluminum and rubber platforms; Fisher Scientific, catalog numbers: 88881122 and 88881123) at 120 rpm speed. On day 9 (maximum day 10), most of the medium was removed and replaced with NPPM without SB-431542 and LDN-193189, but still containing the WNT inhibitor XAV-939. If the medium turned excessively yellow, indicating pH acidification, cookies were redistributed across multiple wells with a maximum of 5 cookies per well to reduce metabolic waste accumulation. On day 20, the medium was switched to Neural Differentiation Medium (NDM) consisting of 50% DMEM/F12 and 50% Neurobasal medium (ThermoFisher Scientific, 21103049) supplemented with N2, GlutaMAX, NEAA, NaPyr, 50 µM β-mercaptoethanol, 2 µg/mL heparin, and insulin (25 µL per 100 mL; Sigma, 19278). On day 30 (maximum day 35), NDM was replaced by Long-Term pro-Survival Medium (LTSM), consisting of Neurobasal medium supplemented with N2, B27 without vitamin A, GlutaMAX, NEAA, NaPyr, 50 µM β-mercaptoethanol, 2 µg/mL heparin, 1% serum (Fetal Bovine Serum; ThermoFisher Scientific, 10270-106; heat-inactivated at 56 °C for 30 minutes, aliquoted and stored at −20 °C), 10 ng/mL BDNF (PeproTech, 450-02), and dissolved Matrigel (final concentration in the medium: 0.1%). Medium was changed every 3–4 days until the desired stage of differentiation. Starting from day 50–60, larger organoids were sliced under a stereomicroscope using a sterile blade (typically once per month), which improved long-term survival and limited the formation of necrotic cores [20]

***Mesoderm differentiation for endothelial organoid (EO) production***

Mesoderm differentiation protocol was adapted from Wimmer et al [21]. Dissociated hiPSC cells were seeded into ultra-low attachment 96-well plates at a density of 4,000 cells/well in mTeSR1 medium supplemented with 5 µM dihydrochloride ROCK inhibitor Y-27632. To facilitate embryoid body (EB) formation, plates were centrifuged at 125 g for 4 minutes to aggregate cells at the bottom of the wells, and further incubated in a humidified atmosphere at 37 °C with 5% $CO_2$. Two days later, the medium was changed and EBs were cultured for an additional 3 days in DMEM/F12 medium supplemented with N2 and B27 (with vitamin A) supplements, 10 µM CHIR99021 (TOCRIS, 4423), and 30 ng/mL BMP4 (ThermoFisher Scientific, PHC9534) to induce differentiation toward the mesodermal lineage. On day 5, the medium was removed and replaced with N2/B27 medium containing 40 ng/mL VEGF-A and 2 µM forskolin (TOCRIS, 1099) for a further 2-day incubation period. To generate vascular organoids (EOs) with blood vessel-like networks, 10–15 mesodermal EBs were seeded onto a solidified Collagen I (Corning, 354236)/Matrigel matrix layer in 12-well plates and fully embedded in the same matrix. (Refer to the original publication for collagen solidification steps using HEPES and NaOH solutions.). As an alternative approach to limit EB adhesion and facilitate following protocol steps for assembloid generation, Collagen/Matrigel coating was performed in 24-well plates containing a round glass coverslip at the bottom of each well. After embedding into Collagen/Matrigel gelatin, pre-warmed StemPro34 SFM (ThermoFisher Scientific, 10639011) complete medium containing 10% FBS (Sigma MERCK, F7524), 40 ng/mL VEGF-A, and 20 ng/mL FGF-2 was added to induce blood vessel differentiation. Vessel sprouting was typically observed after 2–3 days. Medium was partially changed every other day. EOs were used for CO+EO assembloid formation always after day 10 and only when successful vessel sprouting was evident. To recover EOs, a cut-open P1000 pipette tip was used, and forceps were applied to gently remove excess matrix from around the organoids.

***Cortico-endothelial assembloid (CO+EOs) formation***

To form CO+EO assembloids, pairs of one cortical organoid (CO) and one endothelial organoid (EO) were co-cultured in low-adhesion 96-well plates in Long-Term pro-Survival Medium (LTSM) supplemented with 40 ng/mL VEGF-A and 20 ng/mL FGF2. When COs were significantly larger than their endothelial counterparts, they were micro-dissected into smaller fragments using forceps prior to assembloid formation. Medium was partially refreshed daily. Fusion was allowed to proceed for one week before GB cells addition. For GB cells incorporation, one week after the initial co-culture, 10,000 GB cells were added directly on top of the well containing the assembloid. The plate was centrifuged at 300 rpm for 1 minute to promote cell sedimentation and contact with the organoids, followed by a 24-hour incubation to facilitate GB cell integration. The next day, CO+EO assembloids were transferred either onto an orbital shaker or into a SpinΩ mini-rotor to support optimal growth until the desired endpoint.

***Immunostaining on cryostat sections***

Immunostaining on cryostat sections was performed as previously described [20]. Briefly, fixed organoids and assembloids in 4% PFA at 4 °C for 4 hours were dehydrated in 10% sucrose for at least 4 hours to overnight, followed by 25% sucrose (overnight at 4 °C). Sucrose was then substituted with OCT resin (Leica Tissue Freezing Medium, 14020108926; or Cryomatrix™, 6769006), and organoids and assembloids were transferred into embedding molds in clean OCT and stored at −80 °C. Cryostat sections of 12 µm thickness were cut using a Leica cryostat (model: CM3050S), collected on glass slides (ThermoFisher Scientific, Superfrost Plus, J1800AMNZ; or VWR SuperFrost™ Plus, 631-0108), and stored at −80 °C. Prior to immunostaining, sections were dried at room temperature for 10–15 minutes, then washed twice with PBS 1× (10 minutes each) to remove traces of OCT resin. For some antibodies, antigen retrieval was required prior to primary antibody incubation and performed by unmasking in 0.1 M sodium citrate solution (pH 6) at 95 °C for 10 minutes. A 5-minute PBS 1× wash was then performed to remove the unmasking solution, followed by addition of a pre-blocking solution containing PBS 1× with 5% serum (sheep serum, Sigma, S2263-100ML or newborn calf serum, ThermoFisher Scientific, 16010-167) and 0.3% Triton X-100 (Sigma-Aldrich, T8787) for 1 hour at room temperature. For primary antibody incubation (from a minimum of 4 hours at room temperature to overnight at 4 °C), a blocking solution composed of PBS 1× supplemented with 1% serum and 0.1% Triton X-100 was used. Primary antibodies used in this study are listed in Table 1. Alexa Fluor 488, 555, 594, and 647-conjugated anti-mouse, anti-rabbit, anti-rat, anti-goat, anti-guinea pig, or anti-sheep IgG secondary antibodies (ThermoFisher Scientific, all diluted 1:500) were used and incubated from a minimum of 2 hours at room temperature to overnight at 4 °C. The secondary antibody solution also contained Hoechst 33342 (1:1,000; Invitrogen, H3570) for nuclear staining. After final washes (3 times, 10 minutes each in PBS 1×), organoid sections were mounted with a solution composed of 80% glycerol and 2% N-propyl gallate in PBS 1×, covered with glass coverslips, and sealed with nail polish around the edges. Stained sections were stored at −20 °C. Imaging was performed using the Vectra Polaris slide scanner (Akoya Biosciences), followed by artificial intelligence-driven analysis using HALO software (Indica Labs). Alternatively, images were acquired using an Apotome Zeiss system with AxioVision software and exported as TIF files. Confocal imaging was performed on a Zeiss LSM710 confocal microscope using 10×/0.45 NA and 63×/1.4 NA objectives, and images were analyzed using Fiji or Adobe Photoshop.

Supplemental table 1. Antibodies used in immunostaining experiments in this study

| Primary antibody | species | Concentration used | Antigen retrieval | Brand | ref |
|---|---|---|---|---|---|
| CD31 | mouse | 1/1000 | NO | Abcam | Ab9498 |
| VE-Cadherin | Rabbit | 1/2000 | NO | GENTEX | GTX132982 |
| CELF2 | Rabbit | 1/1000 | YES | MERCK | HPA035813 |
| Cleaved Caspase 3 | Rabbit | 1/1000 | YES | Cell signaling | #9661 |
| CLDN5 | Rabbit | 1/1000 | NO | Abcam | AB 131259 |
| OCLN | rabbit | 1/1000 | NO | GENTEX | 114949 |

| | | | | | |
|---|---|---|---|---|---|
| ZO-1 | rabbit | 1/500 | NO | Gentex | 636399 |
| AHNAK | RABBIT | 1/500 ICC; 1/200 WB | NO | INVITROGEN | PA5-53890 |
| SERPING1 | RABBIT | 1/1000 | NO | INVITROGEN | MA5-46954 |
| CHI3L1 | Rabbit | 1/1000 | NO | ABCAM | Ab77528 |
| DCX | Rabbit | | NO | Cell signaling technology | 40619 |
| TUBBI | mouse | 1/10 000 (WB) | NO | SIGMA | T7816 |
| GFP | Chicken | 1/1000 | NO | Abcam | Ab13970 |
| KI67 | Rabbit | 1/1000 | YES | Abcam | Ab15580 |
| MAP2 | Mouse | 1/1000 | NO | Sigma | M4403 |
| OLIG2 | Mouse | 1/1000 | YES | Millipore | MABN50 |
| TUJ1 | Rabbit | 1/1000 | NO | BioLegend | 802001 |

***Immunostaining image analysis on HALO software***

Artificial intelligence analyses were performed using the Random Forest Classifier module in HALO software. The AI was trained on multiple sections and slides to recognize specific tissue areas based on marker expression. The cortex was identified based on MAP2 or TUJ1 expression, the tumor area by GFP, and the endothelial region by the absence of MAP2 and/or the presence of CD31 expression. Once adequately trained, the classifier was saved and applied to additional sections for analysis. HighPlex FL modules (versions 4.2.3 and 4.2.14) were used to detect marker-positive cells. Subsequent analyses were performed using the classifier, as needed.

***Immunostaining and tissue clearing of whole organoids***

The tissue clearing protocol was adapted from a previously described protocol [22]. Fixed organoids were washed with PBS containing 0.05% sodium azide (Sigma; S8032-100G) and stored in PBS-azide at 4 °C until required for immunostaining. A permeabilization step was first performed by incubating the organoids in PBS containing 0.5% CHAPS (Sigma; C9426-5G) for 3 hours at 37 °C. Following this, the permeabilization solution was replaced with a blocking solution consisting of PBS-azide, 0.3% Triton X-100, and 5% newborn calf serum (NBCS), and incubated overnight at room temperature with gentle agitation. The following day, primary antibodies were added at a 1:250 dilution in PBS-azide containing 0.3% Triton X-100 and 5% NBCS, and organoids were incubated for 2 to 3 days at room temperature with gentle agitation. After primary antibody incubation, organoids were washed in PBS-azide containing 0.3% Triton X-100 for a total of 6 hours, with the washing solution changed every hour. Secondary antibodies were diluted 1:500 in PBS-azide containing 0.3% Triton X-100 and 5% NBCS, and incubated with organoids for 24 to 48 hours at room temperature with gentle agitation. Following secondary antibody incubation, washes were performed in the same manner as for primary antibodies. After the final washes, the tissue clearing solution (AKS) was added for at least 24 hours. The AKS clearing solution was composed of 20% DMSO (Thermo Scientific; 20688), 40% 2,2'-Thiodiethanol (TDE) (Sigma; 88561-1L), 20% sorbitol (Thermo Scientific; 036404.36), and 6% Tris base (0.5 M) (VWR; VWRC28808.294), all diluted in distilled water. Organoids were stored in this solution until imaging. On the day of imaging, organoids were embedded in 0.8–0.9% agarose prepared in AKS solution. Imaging was performed using a Light Sheet Fluorescent Microscope system (Lightsheet 7, Zeiss) with 5X objective (Illumination 5X NA 0.1, detection 5X NA 0.16 with Axio Cam 701) and a refractive index of 1.48. Three-dimensional images were reconstructed and analyzed using Imaris software (version 9.6). Organoid and tumor volumes were quantified using the Surface module. Surface classification was selected for subsequent analyses, and surfaces were smoothed by setting the smallest object size to 15 µm. Cell distances from the organoid surface (previously generated) were calculated using the Spot Detection module, with the threshold manually adjusted based on the mean GFP intensity. A color code was applied to enhance visualization.

***Immunostaining of cultured cells***

Cells were smeared on glass coverslips and processed as indicated in the legends. Labeling was performed with antibodies listed in the Supplementary Materials, as previously reported. They were revealed with the appropriate secondary antibody coupled to Alexa Fluor (1:1000). To evaluate the unspecific signal, a control condition without primary antibody and a non-specific antibody was carried out for each antibody. We examined six-to-ten representative fields for each condition. Images were acquired on an inverted AxioObserver-Zeiss microscope equipped with a sCMOS ANDOR Neo camera. The system was controlled by MetaMorph software (Molecular Devices, Sunnyvale, CA, USA). Alternatively, imaging was performed using the Vectra Polaris slide scanner (Akoya Biosciences).

***Western blot***

Cells were rinsed in ice-cold PBS and whole-cell extracts were prepared as previously described (Ravaud et al., Stem Cells, 2015 PMID:25827082 ); briefly, cells or organoids were lysed in cell lysis buffer, sonicated for 10 seconds, and centrifuged at 12,000 g for 10 minutes. Protein extracts (5 to 50 µg, as indicated in figure legends) were resolved by SDS–PAGE under reducing conditions and transferred onto Immobilon-P membranes (Millipore, Molsheim, France). Detection antibodies (listed in the supplementary section) were used according to the manufacturer's instructions, followed by incubation with horseradish peroxidase-conjugated secondary antibodies and signal visualization using an ECL detection kit (Bio-Rad, Marnes-la-Coquette, France). Chemiluminescence was detected using a Fusion FX7 Edge imaging system (Vilber, Marne-la-Vallée, France), and band intensities were quantified using FIJI software.

#### *Gene expression analysis by Real Time RT-PCR*

Total RNA was extracted using the TRI-Reagent kit (Euromedex, Souffelweyersheim, France), and reverse transcription was performed using MMLV reverse transcriptase (Promega, Charbonnières, France) according to the manufacturers' instructions. Real-time PCR was carried out using Power SYBR green PCR Master mix (Applied biosystem, Life technologies, Warrington, UK) or KAPA SYBR FAST (KAPA Biosystems; KK4610) on an ABI Prism One-Step real-time PCR system (Applied Biosystems, Courtaboeuf, France). Gene expression levels were normalized to the housekeeping gene 36B4, and quantification was performed using the comparative Ct (ΔΔCt) method. All primer sequences used in this study are provided in Table 2.

Supplemental table 2. Sequences for the primers used in this study

| Gene | Forward primer | Reverse primer | EFFICIENCY |
|---|---|---|---|
| AHNAK | 5'-CCTGAAAGGGCCTCGGATTT-3' | 5'- GCCTTGAACACCAATGCCTG-3' | 101% |
| BRY | 5'-ATGATGGAGGAACCCGGAGA-3' | 5'-TGGTTCCAGGAAGAAGCCAC -3' | 99% |
| VE-Cadherin | 5'- ATGCGGCTAGGCATAGCATT | 5'- TGTGACTCGGAAGAACTGGC-3' | 97% |
| NRP1 | 5'- GGGGCTCTCACAAGACCTTC -3' | 5'- GATCCTGAATGGGTCCCGTC -3' | 100% |
| VEGFR1 | 5'- acaaggcaagaaaccaagac-3' | 5'- cctcctcttcctcaacatcac-3' | 100% |
| VEGFR2 | 5'- tcagcaggatggcaaagac-3' | 5'- catacttcctcctcctccatac-3' | 98% |
| 36B4 | 5'-CTACAACCCTGAAGAAGTGCTTG-3' | 5'-CAATCTGCAGACAGACACTGG-3' | 102% |

#### *Statistical analysis*

All data were statistically analyzed and graphically represented using Microsoft Office Excel, GraphPad Prism (Version 7.00), or BiostaTGV (INSERM and Sorbonne University, Paris, France). Quantitative data are presented as the mean ± SEM. For cell percentage and number quantification following immunostaining, measurements were performed on at least five sections derived from two to three different organoids, unless otherwise stated. Organoid sections with damaged histology were excluded from further analysis. For cell counting, microscope images were processed using Photoshop or ImageJ software by randomly overlapping fixed-width (100 µm) square boxes on the area of interest (e.g., organoid surface) and quantifying positive cells within the boxes. When calculating percentages over the total cell number, Hoechst+ nuclei were counted unless otherwise specified. Alternatively, marker expression and surface coverage were evaluated using artificial intelligence-driven analysis with HALO software (Indica Labs). Data were analyzed using the Mann–Whitney U-test or two-tailed Student's t-test for comparison between two groups, or by two-way ANOVA for comparisons involving three or more groups. Statistical significance thresholds were set as follows: *$p<0.05$; **$p<0.01$; ***$p<0.001$.

***AHNAK invalidation in patient-derived GB cells***

GB cells were transfected with shRNA Clone sets of three constructs against Human AHNAK nucleoprotein in psi-LVRH1H, including a scrambled control clone (ref # CS-HSH111706-LVRH, Labomics S.A., Nivelles, Belgium) using the jet Prime transfection kit (Polyplus-transfection S.A., Illkirch, France) following the recommendations of the manufacturer. Transfected cells were selected and further grown with hygromycin (50µg/ml). The absence of AHNAK expression was analyzed by western blots as previously described.

**Bulk RNA sequencing and Volcano plot generation**

Bulk RNA sequencing data from shCTRL and shCELF2 GB5 cells), have been previously produced [10], were used to generate volcano plots with the ggplot2 package in R. Differential expression was analyzed between shCELF2 and shCTRL conditions. Gene signatures were obtained from Neftel et al., 2019 [17]
